# Coexistence and cooperation in structured habitats

**DOI:** 10.1101/429605

**Authors:** Lukas Geyrhofer, Naama Brenner

## Abstract

Many natural habitats are structured, which imposes certain environmental conditions on extant populations. Which conditions are important for coexistence of diverse communities, and how social traits in such populations stabilize, have been important ecological and evolutionary questions. We investigate a minimal ecological model of microbial population dynamics, that exhibits crucial features to show coexistence: Populations are repeatedly separated into compartmentalized habitats on a timescale typically longer than growth. In this framework, we consider several scenarios for possible interactions between different strains and their environments, which includes sharing a common nutrient source or expression of public goods that potentially increase population size. Examples for these public good dynamics are collective resistance against antibiotics, and enhanced iron-availability due to pyoverdine. We show that the two features of a long mixing timescale and spatial compartmentalization are already enough to enable coexisting strains. In the case of public goods, stable coexistence immediately entails cooperation.

## I. Introduction

Natural environments are seldom uniform in space and time. For its inhabitants, the spatial and temporal structure of the environment has profound effects on their growth, their interactions and their survival. The effect of structure in space (Nei, 1973; Rainey and Travisano, 1998; Slatkin, 1987; Wright, 1943) and time (Letten *et al.*, 2018; Stewart and Levin, 1973), and the constraints they impose on populations have been studied extensively in the context of population genetics and ecology (Hanski, 1998; Levin, 1976). From these investigations it is generally understood that heterogeneities allow multiple species to coexist (Amarasekare, 2003), with more complex environments usually admitting more diverse compositions of populations. If heterogeneities are hierarchically structured, their impact on the eco-evolutionary population dynamics can be described by multilevel selection (Okasha, 2006; Wilson, 1975): There, fast growth is often favored on lower levels, describing individuals or cells, but the dynamics on all levels can depend on many other factors.

Nowadays, a large effort goes into trying to understand such diverse communities of microbes (Cordero and Polz, 2014). These exist in biofilms, in guts of higher animals, and many other relevant places, where they are important for ecological, economic and medical affairs. For these populations similar issues are of interest: How can these communities be so diverse, how can these groups survive and thrive together, and what role does a structured environment play? Empirical and theoretical answers point towards a few common themes. Diverse populations can interact via coupled metabolisms, where mutualistic cross-feeding (Goldford *et al.*, 2018; Harcombe *et al.*, 2014; Müller *et al.*, 2014) or trade-offs in allocation between multiple resources (Posfai *et al.*, 2017; Taillefumier *et al.*, 2017), both allow for coexistence. Besides these intrinsic mechanisms, spatial structuring and compartmentalization are also found to contribute to diverse microbial populations and their cooperation (Cremer *et al.*, 2011, 2012; Lampert and Tlusty, 2011; Manhart *et al.*, 2018; Matsumura *et al.*, 2016; Melbinger *et al.*, 2010, 2015; Traulsen and Nowak, 2006; Wienand *et al.*, 2015).

The ecological and environmental structuring of microbial communities also allows to address more fundamental problems in evolutionary biology: For example, the evolution of multicellular organisms from single celled ancestors likely required the formation of stable cell collectives engaged in cooperative interactions. The fact that multicellularity evolved multiple times (Grosberg and Strathmann, 2007), hints that the conditions for this are probably not too restrictive. Indeed, experimental studies showed that in yeast multicellular aggregates readily form when environmental conditions impose a selective advantage to groups of cells (Hammerschmidt *et al.*, 2014; Ratcliff *et al.*, 2012; Rose *et al.*, 2018). Understanding the ecological conditions required to form stable collectives can shed light on their evolutionary origin. Some researches have argued that such *ecological scaffolding* can provide the necessary support the evolutionary transition of collectives into new individuals themselves (Black *et al.*, 2018; Rainey *et al.*, 2017).

One natural example of an ecological system with strongly structured environments are tidal cycles on rocky shores (Blaustein and Schwartz, 2001; Dayton, 1975; Sousa, 1979) (see also Fig. 1): High tide dilutes populations into small tidal pools and replenishes nutrients, while at low tide remaining cells utilize these resources to replicate. This cyclic tidal dynamics may be more complex, but its crucial features include the spatial segregation of pools, and a temporal scale determined by the tides, which is long compared to growth. These features can be realized in contemporary laboratory experiments, for instance by enclosing populations in milli- and micro-fluidic droplets (Baraban *et al.*, 2011; Cottinet *et al.*, 2016), which are pooled and then seeded periodically into new droplets with fresh medium.

**Figure 1.**
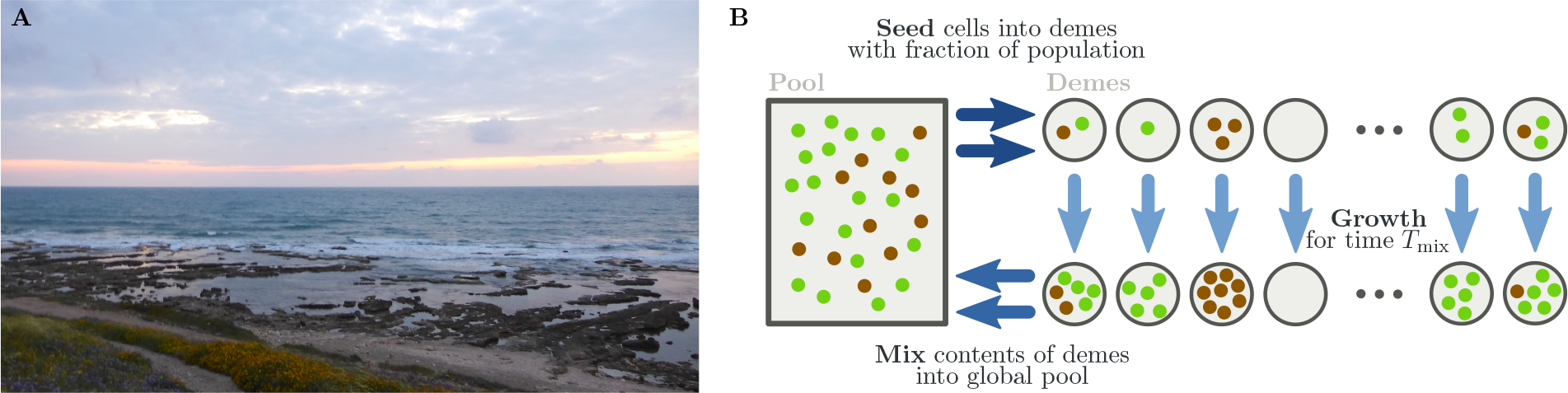
(A) A rocky shore exposed to tidal cycles can represent an example for the dynamics considered in this article. Nutrients are replenished and contents of all small tidal pools are mixed during high tide, while allowing for segregated growth during low tide. Such an environment allows coexistence and cooperation between multiple microbial strains. Pictured is the coastline of Haifa (Israel) near Tel Shikmona. **(B) Schematic depiction of cycles of growth, mixing and reseeding**. Our model of many microbial populations growing in compartmentalized demes can be described by multilevel selection. Two levels are given by the growth dynamics within demes, and the cyclic dynamics of growth, mixing and reseeding on a longer timescale.

In this article, we analyze a minimal model of this environmental structuring, which combines spatial segregation with limited resources and a temporal cycle of mixing and reseeding. We develop a mathematical framework, suitable for microbial populations, which can encompass various interactions between multiple microbial strains. Growth-related processes within a single habitat are described by deterministic differential equations, that include all interactions between strains and interactions with their shared environments. The dynamics on the second, longer timescale of mixing cycles is modeled by a discrete map for the distribution of inoculum compositions, where the seeding of new demes is stochastic.

In this framework, we show that coexistence is a very generic outcome with these two conditions of compartmentalized populations and a long mixing timescale. In particular, we compare a simple growth model with shared resources, to the enzymatic degradation of antibiotic hazards, and to resource extraction via siderophores, where the latter two are examples for public good dynamics. While the specific dynamical interactions are different, coexistence – and thus cooperation via public goods – between strains can be mediated by this spatio-temporal structuring of the environment.

## II. Population Dynamics in Spatio-Temporally Structured Habitats

Our model describes microbial populations growing in a large number of compartmentalized habitats (called demes) for a time much longer than their doubling time. Within a deme these populations feed on a single resource, which can deplete such that growth terminates at a time *T*_depl_. Contents of all demes are mixed after time *T*_mix_ into a common pool. Most of the time, we assume that *T*_mix_ is slightly larger than *T*_depl_, such that resources are depleted, but cell death is not yet an important aspect of the population dynamics. After mixing, the pool is diluted by a factor *d* and cells are again seeded into demes: The cycle starts anew with another round of growth, which is terminated again at *T*_mix_. In Fig. 1B these cycles are illustrated schematically. This cyclic dynamics of growth, mixing and reseeding is qualitatively consistent with tides on a rocky shore, and more quantitatively with distributed millifluidic droplet experiments recently developed for laboratory studies of microbial populations (Baraban *et al.*, 2011; Cottinet *et al.*, 2016).

The two types of dynamics – growth within demes, and the overall cycles – are largely decoupled. In order to separate them also in notation, we always use lowercase letters, like *n_i_*, to indicate quantities in an inoculum. In general, the inoculum is defined as the set of all seeded cells in a deme. An index (usually *i*) denotes strain *i*, and we do not specify the actual number of strains further. For representations in figures we only use two strains. The observables in the inoculum serve as initial conditions for the dynamics within demes, where the explicitly time-dependent quantities are given by uppercase letters, like *N_i_*(*t*). A summary of the notation can be found in Table I.

**Table I.**
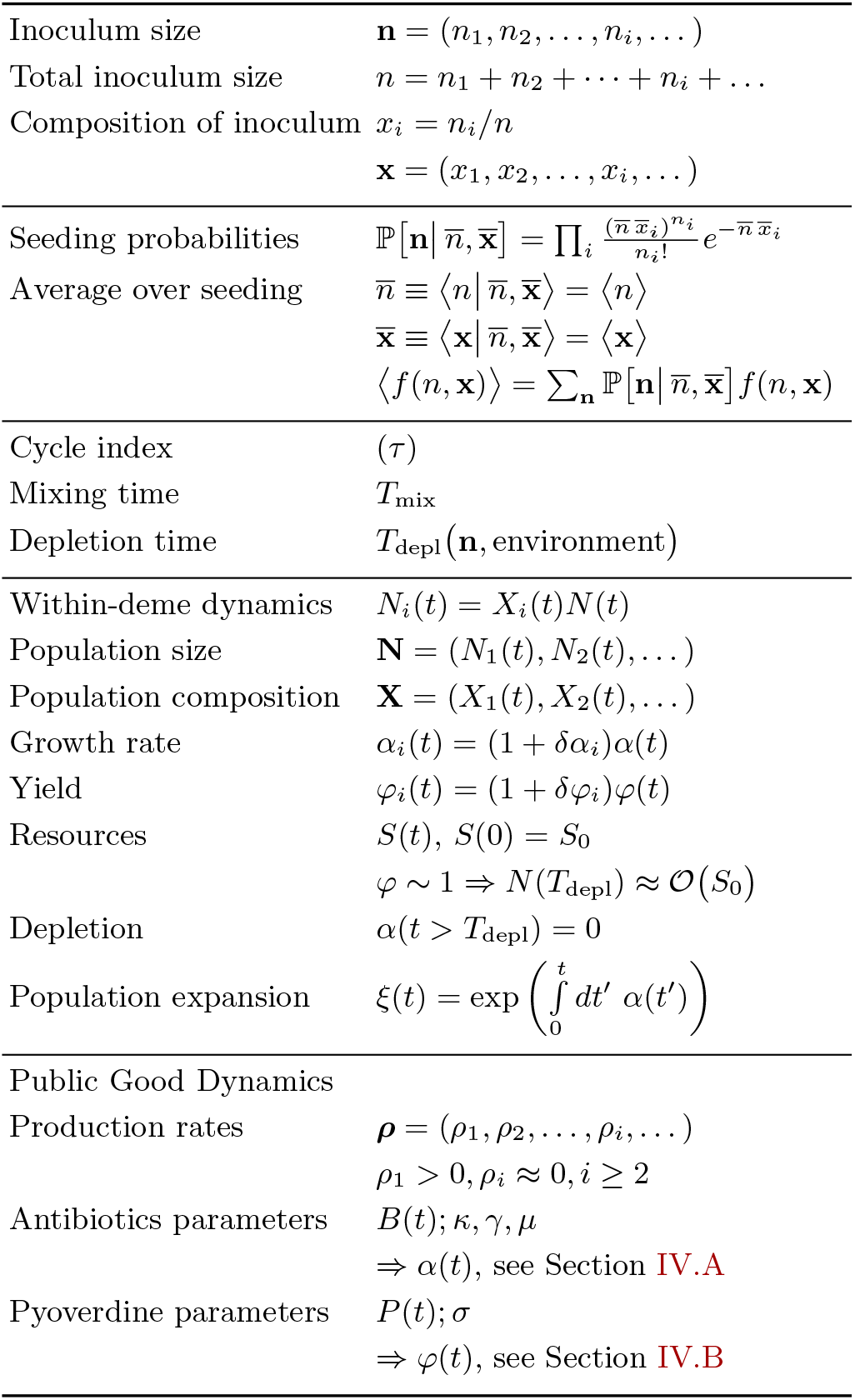
Notation used throughout the main text.

### A. Long-term cycles of mixing and reseeding

For the dynamics of the longer timescale of cycles, only the initial and final numbers of cells in each deme are important. A single deme can be seeded with multiple strains, described by the vector of inoculum sizes **n** = (*n*_1_*, n*_2_,…). The final number of cells for each strain, **N** = (*N*_1_*,N*_2_,…), depends on this inoculum size **n**, and how strains interact during growth. All environmental conditions are assumed to reset to identical values at the time of seeding, which includes replenishing resources. Then, growth can be represented by a deterministic map from initial conditions to the vector of final population sizes at the time *T*_mix_,

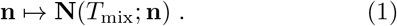

In order to analyze the dynamics over cycles, we focus of the distribution of inoculum sizes over all demes. How this distribution maps to a new distribution of inoculum sizes in the next cycle is sufficient to understand the overall dynamics. This can already be answered using the general form of growth in Eq. (1). Details of these growth processes are specified below, in section II.B.

If the inoculum is small compared to the size of the total pool, we assume that **n** in a single deme follows an independent Poisson distribution for each strain, 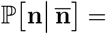 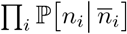 and 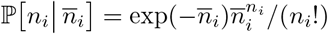. A Poisson distribution is already characterized by its mean 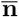, such that we only need to follow the dynamics of these average values. This assumption of a Poisson distribution in the inoculum is consistent with microfluidic experiments, when only few cells are enclosed in a droplet (Bachmann *et al.*, 2013; Baraban *et al.*, 2011; Cottinet *et al.*, 2016). In order to start a new cycle, the pool is diluted by a factor *d* ≪ 1, which indicates the ratio of the average population sizes at the end of the last cycle to the new average inoculum sizes. Then, 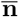 evolves as

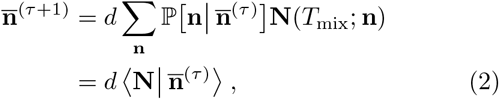
with the index *τ* counting the number of elapsed cycles. The angular brackets used in Eq. (2) denote the averaging over all demes, implied by the global mixing step. In general, this cycle dynamics is a non-linear map from the average inoculum size in the last cycle to the average inoculum size in the new cycle, 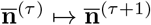. When we want to emphasize this dependence we explicitly write 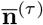 after a vertical line inside the angular brackets. Most of the time, however, we just use e.g. 〈*N_i_*〉 for average final population sizes started from the various seeding compositions.

For the cycle dynamics, all cells are collected from every deme, and during seeding they are diluted and distributed into the same number of demes – so the actual number of demes drops out; It is only important that enough demes exists, that the inoculum size distribution can be sampled sufficiently.

To follow the long-time-evolution of the inoculum, a transformation of variables provides additional insight: we define the total sizes in inoculum and during growth as *n* = Σ*_i_n_i_* and *N* = Σ*_i_N_i_*. In order to track how this population is composed, we introduce the fractions *x_i_* = *n_i_*/*n* and *X_i_* = *N_i_*/*N*, which we summarize again as vectors x and **X**. Then, the mapping for inoculum size dynamics can then be stated as

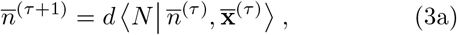

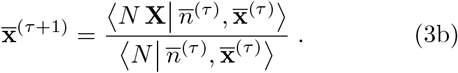

Due to the *global* pooling step we average the total number of cells for each strain 〈*NX_i_*〉, and the total population size 〈*N*〉 *before* taking their ratio. The dynamics of the population composition can be reformulated using the covariance, 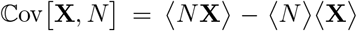, solved for 〈*N****X***〉/〈*N*〉. Then, defining Δ**X**(*T*_mix_; *n*, **x**) ≡ **X**(*T*_mix_; *n*, **x**) − **x** as the change in composition during growth in a single deme, we can subtract 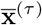 from Eq. (3b), rendering it equivalent to

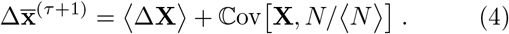

The dynamics in Eq. (4) is analogous to the *Price equation* (Gardner, 2008; Price, 1970), which is one possible framework for multilevel selection (Okasha, 2006). In general, the Price equation describes the dynamics of a quantitative trait (in our case the average composition within demes) due to its inherent transmission bias and how it covaries with fitness (here the final population sizes). In our model two terms have a clear interpretation on which level of selection they act: The average change of the population composition, 〈Δ*X_i_*〉, describes selection among cells within a single deme. Typically, 〈Δ*X_i_*〉 will be positive for strains with larger growth rates. The second term 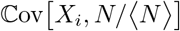 indicates selection among different demes in the pooling and reseeding step, where larger final sizes are favorable at this level of selection over the longer timescale of cycles.

The dynamics in Eq. (4) allows to encode effects that have been termed *Simpson’s paradox* (Blyth, 1972; Simpson, 1951). In general, these Simpson-type effects are counter-intuitive statistical observations that require additional structure of the underlying data. The illustration in Fig. 2 depicts this situation: Local competition shows the opposite outcome from the global dynamics in the frequency dynamics. There, the ‘green’ strain always declines in frequency within each group, 〈Δ*X*green〉 < 0, as it is slower growing. However, a larger initial fraction of the ‘green’ strain correlates with larger final sizes, 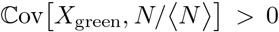. If this correlation is strong enough, then due to the global pooling, the ‘green’ strain can increase in frequency over cycles, 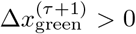. Exactly this mechanism is important for coexistence (or even fixation) of costly traits in the population.

**Figure 2.**
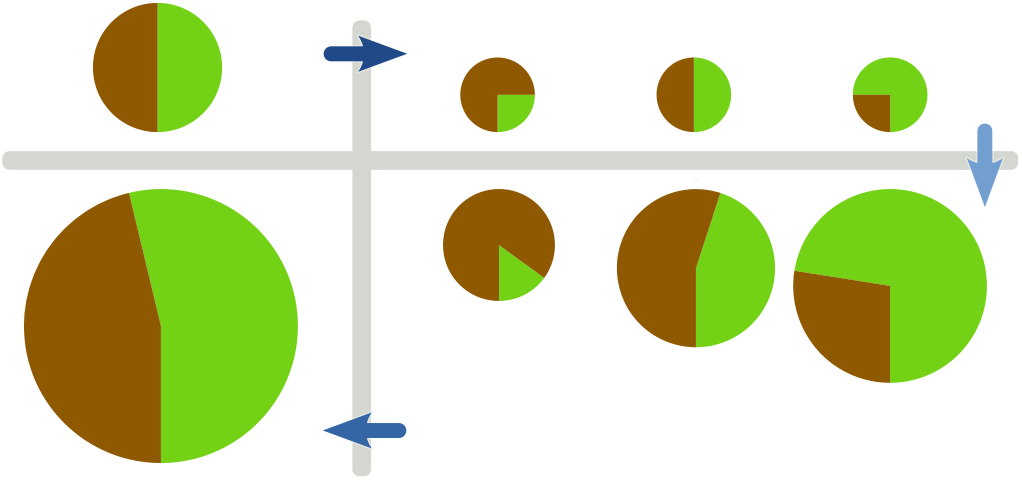
‘Simpson’s paradox’ visualized. If a strain decreases in frequency in direct competition (even in *each* group, as shown on the right), its overall frequency can still increase in the total pooled population (on the left), when the composition positively correlates with final group sizes. In order to observe this effect, two conditions are required: variation in initial conditions, and large enough differences in final sizes. Both of these will occur readily in our model: small inoculum sizes will induce different composition of initial conditions, while different population compositions result in variation in final size from either differences in resource efficiency, or other social traits.

As stated in Eq. (1), our model does not include stochastic effects for the growth processes within a single deme itself. However, the seeding process with its distribution of inoculum sizes provides already a stochastic component in the overall dynamics. This stochastic seeding is crucial for Simpson-type effects to occur: If all populations are seeded with identical initial conditions, the distinction between different demes does not provide any additional structure. Mathematically, the covariance term in Eq. (4) would vanish without variation in seeding.

### B. Growth dynamics within demes

Within a single deme, we consider the growth dynamics of multiple strains *N_i_*(*t*), and the time-evolution of the resource concentration *S*(*t*):

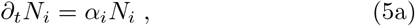

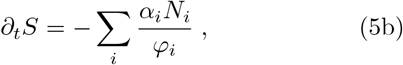
where different strains *i* are characterized by their growth rates *α_i_* and yields *φ_i_*. In general, these two parameters *α_i_*(*t*) and *φ_i_*(*t*) can be time-dependent or influenced by additional observables, that are not yet contained in Eqs. (5). Such explicit time-dependence can also imply a dependence on *N_i_*(*t*) or *S*(*t*). Indeed, we assume that *α_i_*(*t*) = 0, when resources are depleted within a deme, *S*(*t*) = 0 for *t* > *T*_depl_. This process of depletion is explicitly modeled by Eq. (5b), where the initial amount of resources *S*_0_ decreases with each newly grown cell, using yield *φ_i_* as conversion factor between cell numbers and resource concentration. Thus, the decreasing resource concentrations can be seen as a timer to stop the growth dynamics.

Similar to the dynamics over cycles, we use total population size *N*(*t*) and fractions *X_i_*(*t*) of different strains, which transforms Eqs. (5) into

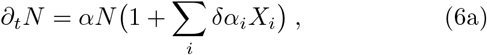

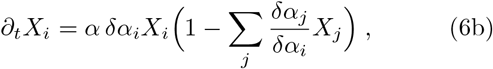

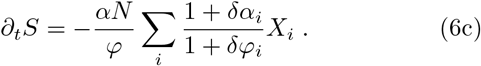

Here, we separated the average growth rate *α* and average yield *φ* from deviations of individual strains, *α_i_* = (1 + *δα_i_*)*α* and *φ_i_* = (1+ *δφ_i_*)*φ*. These averages *α* and *φ* are defined as arithmetic mean over all strains, and are independent of the individual compositions within demes. Eq. (6b) describes the frequency changes in a multi-species Lotka-Volterra model, also known as replicator dynamics, which has been studied extensively (Hofbauer and Sigmund, 1998; Murray, 1989; Nowak, 2006). For us, however, the dynamics of the population size *N* is important as well, because interactions between microbial strains do not only change the frequency dynamics, but also how frequency dynamics couples to the population size dynamics. The covariance term in Eq. (4) indicates exactly this fact.

A simplifying assumption, that will help to solve Eqs. (6), is to allow time-dependent *averages* of growth rate *α*(*t*) and yield *φ*(*t*), while differences between strains, *δα_i_* and *δφ_i_*, are constant. This separation is useful for microbial populations, where differences are likely small. With the average growth rate *α*(*t*), we can define the *average expansion factor* of a population as

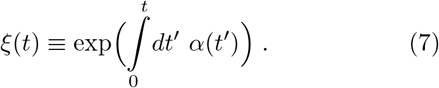

For the special case of constant growth rate, *α*(*t*) = *α*, this reduces to *ξ*(*t*) = exp(*αt*). Using *ξ*, the solutions to total size *N* and population fractions *X_i_* in Eqs. (6) can be stated as

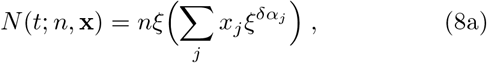

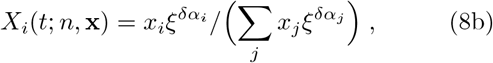
with *n* and **x** the initial conditions in a given deme. The time-dependence in these solutions enters only via the integration limit in *ξ*(*t*). Besides separating time dependence from initial conditions, another advantage of definition (7) is that interactions between strains – such as public good dynamics – will manifest as changes in *ξ* as well, as we explain below.

In the mapping over cycles (3), we are interested in solutions of the within-deme dynamics at the time of mixing *T*_mix_. One step towards obtaining these values is computing the expansion factor at the time of depletion, *ξ*(*T*_depl_), which can be derived by integrating resource use, Eq. (6c), from *S*(0) = *S*_0_ to *S*(*T*_depl_) = 0,

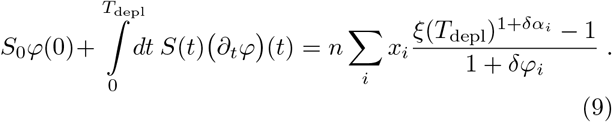

For time-constant yields *φ*(*t*) = *φ*, and a sufficiently large amount of resources, such that *ξ* ≫ 1, this relation simplifies: The integral with its explicit time-dependence drops, and *T*_depl_ only enters via the final value of the expansion factor *ξ*(*T*_depl_). Consequently, Eq. (9) reduces to

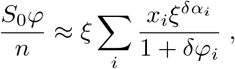
which we can approximately solve as

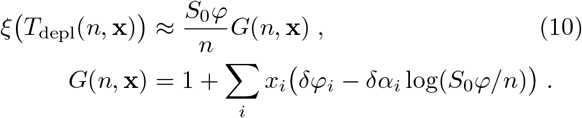

For small differences between the strains, *δα_i_,δφ_i_* 1, we have *G* ≈ 1 and thus to leading order *ξ* ≈ *S*_0_*φ/n*. For the resource consumption dynamics, *ξ*(*T*_depl_) including the corrections *G*(*n*, **x**) can be interpreted as a (non-linear, but monotonic) transformation of the depletion time *T*_depl_: With that context, in demes with increased yield, Σ*_i_x_i_δφ_i_* > 0, strains will grow longer (larger *ξ*), while in demes with fast growth, Σ*_i_x_i_δα_i_* > 0, the time to depletion decreases (smaller *ξ*).

For the cycle dynamics in Eqs. (3), we still need the value of *ξ*(*T*_mix_) instead of *ξ*(*T*_depl_). However, we assume that strains stop growing when resources are depleted, and *α*(*t*) = 0 for *t* > *T*_depl_. Then, *ξ* stays constant for depleted resources. In the opposite case, *T*_mix_ < *T*_depl_, not all resources are used up. We show in Appendix A that depleted resources are actually a necessary condition for the stable growth of a strain in the cycle dynamics. Each strain that does not deplete all nutrients when grown alone will ultimately go extinct. Thus, the relevant case is faster depletion than mixing, *T*_depl_ < *T*_mix_, such that *ξ*(*T*_mix_) = *ξ*(*T*_depl_), and we will use the expression (7) as *ξ*(*T*_mix_), as long as the within-demes leads to depleted nutrients.

## III. Isoclines of the Cycle Mapping

Before analyzing how public goods shape interactions within demes, we continue building the framework with the resource consumption dynamics. With both the stochastic seeding in Eq. (3), and the solutions for population size trajectories, Eq. (8), we now combine these two approaches. In particular, we will focus on the structure of isoclines of the cycle mapping. Isoclines are curves in phase space – spanned by the coordinates of the average inoculum 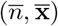 – that indicate conditions where one of these variables stays constant upon evaluating the cycle map, while other variables can change. Each variable in phase space features an isocline, which separates regions it increases, from regions where it decreases over cycles. As our mapping is non-linear, we sometimes find multiple (even non-connected) branches of isoclines. Inspecting all isoclines simultaneously allows to qualitatively map how trajectories evolve over cycles: Fixed points of the mapping exists at the intersection of *all* isoclines, where none of the variables changes.

To obtain these isoclines, we need to evaluate the relations

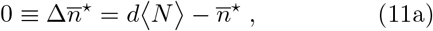
for the total population size and

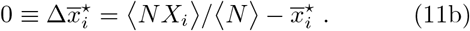
for each of the population fractions. In order to specify the isocline, we use a superscript * to mark the variable that is constant over one cycle.

As we will see below, two main parameters emerge, that determine the position of these isoclines in phase space, while leaving their general shape invariant. Explicitly, the position of the population size isocline, 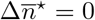, is largely determined by the dilution rate *d*. In contrast, the population composition isoclines, 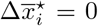, are mostly determined by growth rate differences *δα_i_*. An important observation is that these parameters do have negligible influence on the respective other isocline. Thus, we argue that changing dilution rate *d* will often allow to find intersecting isoclines and generate a coexistence fixed point, where multiple strains are present and stable over cycles.

Illustrations of our results will only use two strains (*i* = 1, 2). Then, the dynamics can be specified with only the total inoculum size, 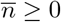, and the fraction of the first strain, 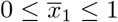. Isoclines will be depicted using *blue* lines for the total inoculum size isocline, and *orange* lines for the fraction isocline. Examples are shown throughout all Figs. 3, 5, 6, 7, 10 and 12. In all figures, we also always assume that strain 1 is slower growing, *δα*_1_ < 0. For the resource consumption dynamics, a trade-off for being more efficient in resource use exists, *δφ*_1_ > 0. Algorithms for computing the exact values of these relations in phase plane (as shown in figures) are presented in Appendix D.

**Figure 3.**
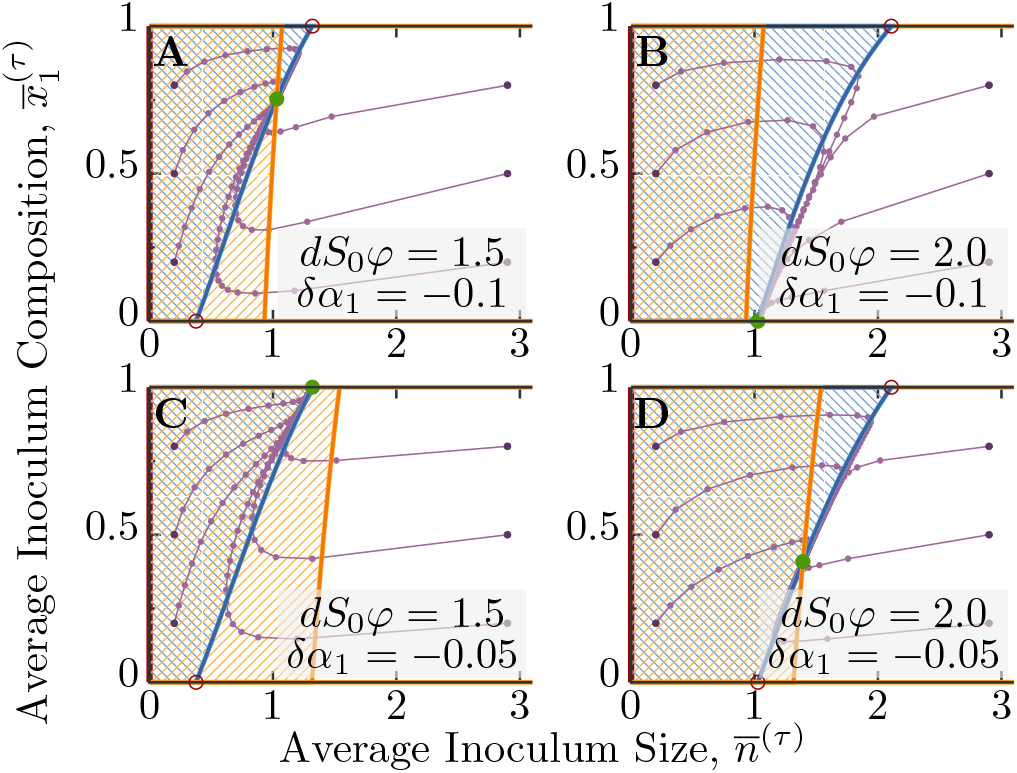
Dynamics of inoculum over multiple cycles with resource consumption. Trajectories of average inoculum size *n* and average inoculum composition 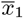 are displayed in purple, where connected dots indicate one cycle of growth, mixing and reseeding. Dark purple points indicate starting points for these trajectories, which are followed for 50 cycles, and can end in stable fixed points (full green points). Empty red points are unstable fixed points, which also includes the whole 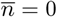 axis. Isoclines for total population size, 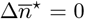, are shown in blue, while isoclines for the population composition, 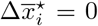, are shown in orange. Hatched regions indicate inocula where either size 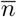 or frequency 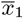 increases over one cycle, while outside the respective parameter decreases. The four panels explore the effect of changing dilution rate *d* and growth rate differences *δα*_1_. Horizontally *δα*_1_ is constant, while vertically *d* is constant. Parameters not stated in panels are *δφ*_1_ = 0.2, *αT*_mix_ = 24.

### A. Population size isocline

The isocline for the total inoculum size, Eq. (11a), separates regions in phase space where inoculum size increases over one cycle, 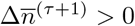, from regions where it decreases, 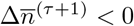. It can be computed by inserting *N*(*T*_mix_; *n*, **x**), Eqs. (8a), into the mapping for *n*(*τ* +1), Eq. (3a), and expanding this expression up to linear order in *δα_i_*, which corresponds to a weak selection limit. As a further step in its derivation, we use the approximation of *ξ* for resource consumption, Eq. (10), which cancels linear terms in *δα_i_* and introduces corrections in yield differences *δφ_i_*. Explicitly, these two steps are given by

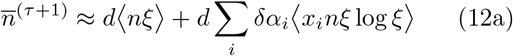

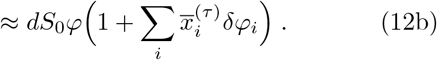

This latter expression only depends on 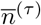 in the neglected sub-leading terms, which implies a few consequences: First, without any dependence on 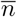, Eq. (12b) is already an approximation for the isocline 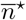 itself, as can be observed when comparing it to Eqs. (11a). Moreover, the average inoculum size is supposed to reach the value determined by Eq. (12) already after a single cycle. In Fig. 3 we show that deviations from this single step approximation occur for small inoculum sizes, which also appear when the linear expansion in *δα_i_* starts to break down. For large inoculum sizes, however, a single step is often sufficient to describe the cycle dynamics for the total inoculum size, as shown for the extended dynamics in Figs. 5 and 7.

From relation (12) we can recognize the main parameter that influences the position of the dilution line is given by the product of dilution rate *d*, the initial substrate concentration *S*_0_ and yield *φ*, as 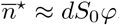. In general, we expect that the dilution rate *d* is the parameter most easily adjusted in experiments. Substrate concentration can also be changed, but this has other implications on the dynamics as well: For instance, depletion time will change, and thus the inequality *T*_depl_ < *T*_mix_ needs to be checked again. Furthermore, growth processes can depend on absolute population sizes. Average yield *φ* just determines how many cells can grow from one unit of resources *S*. It can be chosen 1, such that *S* is measured in the number of potentially growing cells.

Another observation in Eq. (12) is that changes in *δα_i_* only play a minor role in determining the position of the total size isocline. In fact, corrections are only second order 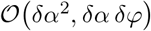, as seen in the computation going from (12a) to (12b). Numerical examples corroborate this observation, shown in Fig. 3, where the blue lines of population size isoclines are almost identical for the two sets of panels AC and BD.

### B. Population composition isocline and Simpson-type effects

The population composition isocline is relevant for treating Simpson-type effects (see Fig. 2): It is the boundary line between regions where frequency of the strain *i* increases, 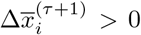, from regions where it decreases, 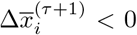. Thus, for strain 1 with *δα*_1_ < 0, we find these Simpson-type effects in the whole region in phase space where the average frequency increases over a single cycle. In addition, both boundaries, 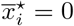 and 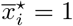, are part of the multi-branched composition isocline: We do not consider additional processes, that would allow for switching between strains or the generation of new strains. Thus, if the dynamics starts on these boundaries it will remain there.

The more interesting cases are branches not on these boundaries: If the isocline cuts through the phase plane for 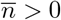, the dynamics potentially exhibits a coexistence fixed point, when the dilution rate *d* is adjusted such that both isoclines intersect. We can derive its condition by inserting the within-deme solutions into the cycle mapping: Assuming again weak selection, we then expand the expressions for frequency change 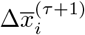, Eq. (4), up to first order in growth rate differences *δα_i_*,

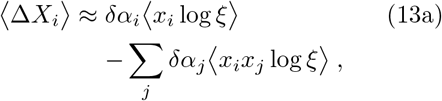

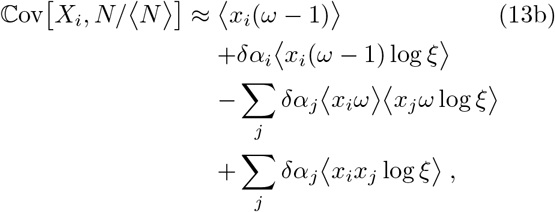
where *ω* = *nξ*/〈*nξ*〉 is a weighting factor describing deviations from the expected final population size.

Similar to before, we can insert the expansion factor *ξ* for resource consumption, which also depends on differences in growth rates and yields, *δα_i_* and *δφ_i_*. Keeping only first order terms in both these differences, we see that two expressions above turn into

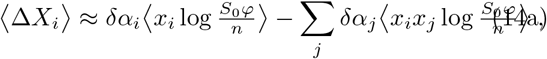

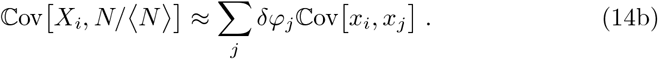

For only two strains, we can explicitly evaluate the averages over the Poissonian inoculum distribution, as we show in Appendix A.3. There, we approximate the composition isocline to leading order as

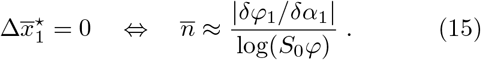

A dependence on the actual fractions 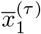 will only appear in higher orders of *δα*_1_ and *δφ*_1_, which is consistent with the very straight (orange) lines depicted in Fig. 3. As a general result, we see that decreasing growth rate differences, will push this isocline to larger inoculum sizes, as can be discerned from Fig. 3 where panels B and D have a smaller value of the difference. This strong dependence on *δα_i_* of the composition isocline is in stark contrast to the position of the population size isocline, where the value of *δα_i_* only appears in higher orders. The dilution rate *d* plays no role in the position of the composition isocline, as it cancels already when calculating the ratio in the cycle mapping, see Eq. (3b).

### C. Coexistence fixed points and their stability

In the last section we worked out the main parameters determining the position of the two isoclines in the resource consumption dynamics: The position of the population size line depends largely on the dilution rate *d*. Deviations from a vertical line in phase space, ie. constant in 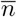, appear due to linearly weighted differences in yield *δφ_i_*, see Eq. (12b). In contrast, the position of the composition isocline strongly depends on growth rate differences. Its shape, however, is given by a constant 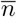, and we find no deviations in 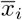 up to linear order in the differences *δα_i_* and *δφ_i_*. We find a coexistence fixed point, when these two lines intersect.

In order to understand the stability of a coexistence fixed point, it is important to examine the stability of the single strain fixed points first. Those exist at the boundaries 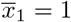 and 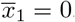, which are both part of the multi-branched composition isocline. Fig. 3 illustrates this situation. The intersections of the population size isocline, Eq. (12b), with these two boundaries determines these two single strain fixed points at 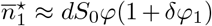 and 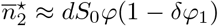. These two expressions can be obtained by evaluating the approximation for the isocline at either *x*_1_ = 1 or *x*_1_ = 0, where the isocline itself linearly interpolates between these two point. The approximations derived for both isoclines, size and composition, show that to first order the population size isocline exhibits a larger tilt, since the composition isocline is almost a straight vertical line. Then, for positive *δφ*_1_ > 0 we can choose a value of the dilution rate *d*, that the larger 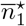 appears in a region where the the fraction *x*_1_ is supposed to decrease, while 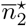 appears in a region where *x*_1_ is increasing (Fig. 3AD): Both single strain fixed points are unstable with respect to invasion by the other strain. Consequently, the intersection in the center of the phase plane, which indicates a coexistence fixed point, will be stable. Fig. 4 depicts these coexistence regions obtained for two strains. For a given dilution rate *d*, coexistence is possible when a trade-off between growth rate differences and yield differences exist. Parameter combinations where where we find this outcome are illustrated in different shades of brown for varying dilution rates. In general, a larger inoculum size (larger *d*) requires a larger yield difference for coexistence. In Appendix A.2 we extend this result, to show coexistence between multiple strains is also possible, when in *each pair* of strains both single strain fixed points are unstable against invasion of the other strain.

**Figure 4.**
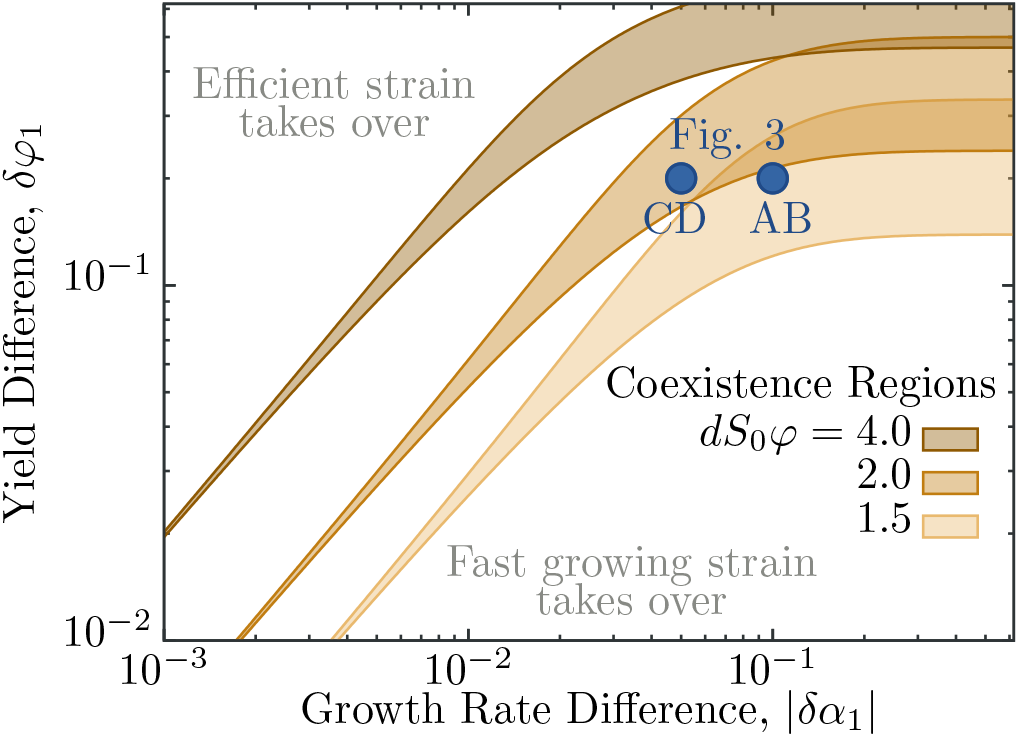
Coexistence regions for resource consumption dynamics within demes. Two strains can coexist in the dynamics over cycles, if they exhibit a trade-off between fast growth and high yield: Shaded regions indicate stable coexistence, outside this region the one of the strains will take over. Here, strain 1 is slower growing, *δα*_1_ < 0, but more efficient in its resource use, *δφ*_1_ > 0. If average number of cells in the inoculum (≈ *dS*_0_*φ*) increases, the efficient strain is less likely to be seeded into demes alone and consequently requires a much larger yield to generate the same number of cells to contribute to the pool at the time of mixing. We observe this effect for large growth rate differences (on the right), where the value of the growth rate difference starts to become irrelevant and final sizes of the slower growing strain in shared demes become negligible. For the parameters *δα*_1_ and *δφ*_1_ indicated by blue dots, we show trajectories in Fig. 3. Depending on the dilution rate *d*, these parameters are either inside or outside the coexistence region.

In natural populations, we also expect that dilution rate can fluctuate, and reach values that allow for coexistence. Fluctuations have also been identified to enable multiple strains to coexist in a similar setting (Ernebjerg and Kishony, 2011).

The previous reasoning, and in particular approximations for isoclines, works well for small *δα_i_* and *δφ_i_*. If growth rate differences become large, however, we can use a different argument that could explain the coexistence observed in the upper right corner of Fig. 4. There, the coexistence region starts to become independent of *δα*_1_, and all demes shared between the two strains will typically end up with almost only the faster growing strain, as it is outgrowing the more efficient strain with a larger yield, *δφ*_1_ > 0. Only if this efficient strain is seeded alone, it can grow to the large final sizes implied by the larger yield. These two effects need to balance for coexistence: The increased final population sizes in demes with only the efficient strain needs to make up for the lower probability of seeding the efficient strain alone. With a Poisson distribution for seeding, this probability is given by 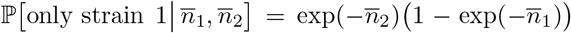. Thus, depending on bottleneck sizes in each cycle, the efficient strain is exponentially suppressed in the Poisson seeding: This coexistence mechanism only works for small inoculum sizes, otherwise it would require unrealistically large *δφ*_1_.

## IV. Public Good Interactions Within Demes

Public goods are environmental compounds that promote or enhance growth, and are available to all cells within a shared environment. Often, public goods need to be actively produced and can generate additional ecological interactions with the environment, which can lead to more complex trajectories *N_i_*(*t*). Here, we analyze two classes of public good interactions: One is collective resistance to antibiotics, mediated by secretion of an antibiotics-hydrolyzing enzyme. The second class of interaction involves active extraction of a resource from the environment, such as iron-chelation by extracellular pyoverdine. These two extensions of the resource consumption model provide examples for a time-dependent growth rate *α*(*t*) in the case with antibiotics, and a time-dependent yield *φ*(*t*) generated by the interactions with pyoverdine. In order to couple these dynamics to our spatio-temporal model, we add an additional dynamical observable to the within-deme system, Eqs. (5), which then influences either *α*(*t*) or *φ*(*t*).

In our formalism, changes in the growth process generate deviations on how *ξ* depends on the inoculum and other environmental conditions. Thus, after stating these additional dynamical observables, we derive its influence on *ξ*(*n*, **x**) and analyze isoclines of the ensuing cycle mapping.

### A. Active reduction of antibiotic hazards

Antibiotic resistance, provided by extracellular enzymes, can be considered a prime example for a public good: After the enzyme has been produced and excreted by a cell, its surrounding population can benefit from reduced antibiotic concentrations. In the following, we consider a scenario where an antibiotic is supplemented with a concentration *B*(0) = *B*_0_ in all demes at each seeding step. If at least one strain is present, that is able to significantly reduce this concentration over time, the overall population may be able to grow again. This reduction of the antibiotic concentration can lead to cross-protection of other strains in the same environment, which has been found experimentally (Domingues *et al.*, 2017; Nicoloff and Andersson, 2015). We assume that resistance to the antibiotics comes at a cost in terms of growth rate, as has been described directly in (Melnyk *et al.*, 2015) and implied indirectly in the results of (Chuang *et al.*, 2009). Moreover, it has been reported that the effectiveness to treat bacterial populations with antibiotics depends on the initial number of present cells (Artemova *et al.*, 2015; Jepson *et al.*, 2016; Tan *et al.*, 2012; Udekwu *et al.*, 2009), especially if the inoculum is small. Exactly this *inoculum effect* will provide the variation in final sizes to generate a large covariance term in the change of composition, such that slower growing, producing strain will coexist or even fixate in the population.

Several observations have been made over the last few decades, on which we base our model: First, growth rate of microbes in the presence of antibiotics is proportional to growth rate in absence of antibiotics (Lee *et al.*, 2018; Tuomanen *et al.*, 1986). Moreover, it is reasonable to assume that the death rate is bounded by a maximal value. Thus, a common model for the growth rate of microbial populations exposed to antibiotics is a sigmoidal Hill-function (Regoes *et al.*, 2004),

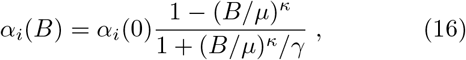
where the base growth rate can vary between strains, *α_i_*(0) = (1 + *δα_i_*)*α*; *B/μ* = 1 indicates the ‘minimal inhibitory concentration’, where growth rate crosses zero and becomes a death rate; *γ* determines the maximal death rate for large concentrations *B, α_i_*(*B → ∞*) = −*α_i_*(0)*γ*; and finally, *κ* indicates the steepness of the transition between growth and death rate around concentrations *B*/*μ* ≈ 1, with large *κ* implying an almost step-like switch from unhindered growth to rapid death, and small *κ* a more gradual transition.

In addition to this effect of antibiotics on growth rate, we explicitly consider the active reduction of antibiotic concentration in a deme,

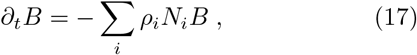
where *ρ_i_* characterizes the rate of resistance of strain *i*, incorporating both expression rate of active enzymes and efficiency of the degradation reaction. This antibiotic concentration *B*(*t*) evolves simultaneously with the dynamics of the population sizes *N_i_*(*t*) and the resource concentration *S*(*t*) in Eqs. (5) in each deme.

Coupling these interactions – how antibiotic concentration changes the overall growth rates in a deme, Eq. (16), and how the antibiotic concentration is reduced, Eq. (17) – generates a race between two processes. Either the amount of antibiotics is large enough to kill all cells within a single deme – or microbes can reduce the concentration below *B*(*t*)/*μ* < 1 before they go extinct, after which the population starts recovering. In this setting, resistance and recovery is mostly a dynamical effect. If we furthermore assume *T*_mix_ long enough, such that recovering populations deplete all nutrients, we will find either a fully grown population or no cells at all. Then, the expansion factor can be stated as

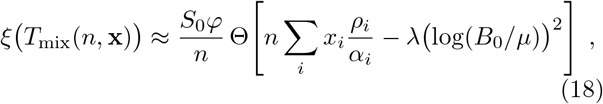
neglecting the sub-leading terms of orders *δα_i_* and *δφ_i_*, see Eq. (10), which would give the multiplicative correction *G*(*n*, **x**). The Heaviside-Theta function Θ indicates the allor-nothing effect on the growth of microbes, which is the main change in *ξ* (since Θ(*y*) = 1 for *y* ≥ 0 and Θ(*y*) = 0 for *y* < 0). Moreover, *λ* = *κγ*/(1 + *γ*) summarizes parameters of the interactions between antibiotic and microbes. Details on the derivation of Eq. (18) can be found in Appendix B.1.

In general, the approximation in Eq. (18) is valid for *B*_0_/*μ* > 1, where cells need to reduce antibiotics, and would die without fast enough reduction. If *B*_0_/*μ* < 1, all populations can already grow at the beginning, and antibiotic reduction only speeds up the growth process. This speed up can generate a large enough differences in final sizes, which in turn can lead to a large enough covariance 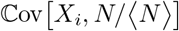 for the producing strain to increase in frequency. Nevertheless, the opposite outcome of the non-producing strain outgrowing the producing strain is possible as well. Details depends on comparing the mixing time *T*_mix_ to the slowed down growth with only few producing cells that lead to a long *T*_depl_. Thus, restrictions on mixing times *T*_mix_ that allow coexistence are very stringent for *B*_0_/*μ* < 1.

#### Analysis of isoclines

The dynamics of antibiotic reduction can be analyzed in the same way as before, via isoclines of the cycle mapping, Eqs. (11). For the following, we only consider a single producing strain, *ρ*_1_ > 0, and one non-producing strain *ρ*_2_ = 0. The cost of production demands *δα*_1_ < 0. Then, we expect only populations to grow that have enough producing cells in the inoculum, with the threshold 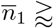 (*λα*_1_/*ρ*_1_)(log *B*_0_/*μ*)^2^ derived from the argument of Θ in Eq. (18). In the cycle mapping, the Poisson distribution in the seeding step smears out this hard cutoff for the resulting final sizes.

At first, we check the limiting case of only a single strain, which produces the public good. Despite the smearing of final sizes, the cycle map of this single strain increases sharply around the threshold of enough producers. This steep increase exhibits either no or two (non-zero) fixed points, depending on the dilution rate *d*. The critical value for *d* can be derived by requiring that the inoculum size determined by resource limitation, Eq. (12b), is larger than than the inoculum size for successful antibiotic reduction. Explicitly, we have 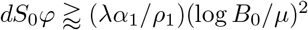. If two fixed points exist, then the larger one is stable (from resource limitation), while the lower fixed point (from antibiotic reduction) acts as boundary for the basin of attraction to extinction. More details can be found in Appendix B.

For two strains, the population size isocline will intersect the 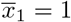 axis at exactly these two values for 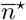, as long as *d* is large enough to fulfill the condition in the last paragraph, see Fig. 5. From the stable fixed point (on the right, with larger inoculum sizes) it extends down to lower 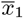 values along the condition derived before for resource consumption, Eq. (12). Without any differences in yield, *δφ*_1_ = 0, it is a straight line. The other condition, limited by antibiotic threat, extends along 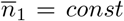, and thus the isocline will scale with 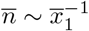. Both parts of the isocline will meet, when 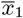 becomes small enough and the isoclines connects these two descriptions.

The population composition isocline, 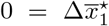, is determined by the balance of the local losses in frequency and the coupling to population size changes, 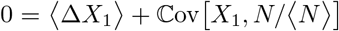 As we assume *δα*_1_ < 0 for the producing strain, the first term is usually negative. Inspecting the expansion in growth rate differences, see Eq. (13), we observe that one term does not depend on *δα*_1_ (contrary to the resource consumption before, factors of *δα_i_* and *δφ_i_* in *ξ* are negligible for antibiotic reduction). This term, 〈*x*_1_(*ω* − 1)〉 is responsible for large values in the covariance, required for coexistence: Inserting *ξ* for antibiotics, 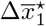 can be written as

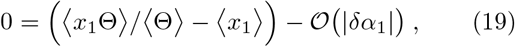
where Θ abbreviates the full expression of the threshold in *ξ*. The first bracket indicates the difference between the average inoculum fraction of surviving populations and the average fraction in the inoculum itself. This term is expected positive, as surviving populations likely have a larger fraction of the producing strain. However, this difference decreases with increasing inoculum sizes, as then more populations survive, and the two averages become closer. Numerically, we find an exponential decay in 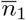, until 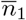 reaches its threshold value in the argument of Θ. For 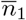 larger than this threshold, the difference decays algebraically with an exponent close to −1. The dependence on the non-producer inoculum size 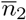 is weak. Thus, 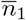 fully specifies the shape of 〈*x*_1_(*ω* − 1)〉 across the whole phase plane 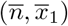. To compute the isocline, the decay of this (positive) expression needs to be balanced by terms of order 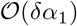; due to this coefficient *δα*_1_ their magnitude varies a lot less. Hence, for the isocline we seek curves where the first term in Eq. (19) is almost constant, which happens along 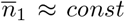. Balancing terms we approximate the scaling of the isocline as

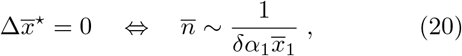
which holds as long as enough cells of both strains are in the inoculum.

If the initial fraction of the producing population is large 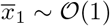, then during the time required to reach *B*(*t*)/*μ* < 1 the non-producing strain can go extinct if it started from too small inoculum sizes. This effect is visible in Fig. 5, when the (orange) composition isocline asymptotically approaches the 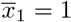 axis.

The influence of the two other parameters of initial antibiotic concentration *B*_0_/*μ* and the resistance rate *ρ*_1_ can also be discerned by analyzing the threshold in the expansion factor, Eq. (18). Details can be found in Appendix B.2.

**Figure 5.**
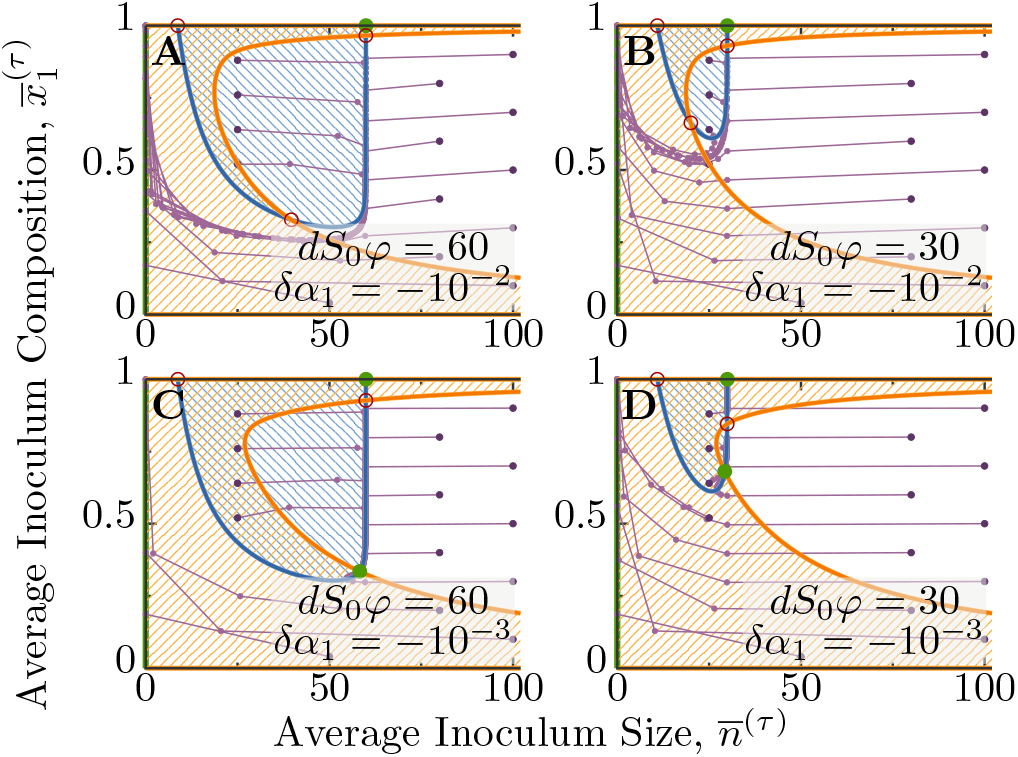
Dynamics of inoculum over multiple cycles with a collectively reduced antibiotic hazard. Trajectories in inoculum size 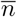 and inoculum fraction 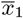 are shown in purple, starting at dark purple points. In general, strain 1 grows slower, *δα*_1_ < 0, and reduces the antibiotic concentration, *ρ*_1_ > 0, while strain 2 does not contribute to the degradation, *ρ*_2_ = 0. During the initial time of within-deme dynamics the dying population needs to reduce the antibiotic concentration before all cells can grow again. For the dynamics over cycles, the blue and orange lines indicate the isoclines for total inoculum size and inoculum composition. Blue hatched areas indicate an increase of the average inoculum size 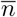 and orange hatched regions depict an increase in the average population composition 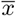, with isoclines shown as solid lines. The four panels show two values of growth rate differences, *δα*_1_ = −10^−2^ (A,B) and *δα*_1_ = −10^−3^ (C,D). Dilution rates (indicated by average inoculum sizes for growing populations) are *dS*_0_*φ* = 60 (A,C) and *dS*_0_*φ* = 30 (B,D). All panels exhibit the same initial antibiotic concentration *B*_0_/*μ* = 1.25, and *ρ*_1_/*α* = 5 · 10^−3^.

Similar to the resource consumption, we again established the dilution rate *d* as the main parameter for the position of the population size isocline: For coexistence, the relevant part of this isocline is identical to the derivation before. In contrast, the position of the composition isocline depends strongly on small changes in *δα*_1_. Again, changing any of these two parameters barely influences the shape of the respective other isocline, as we show in Fig. 6. In this figure, we sweep through 2 orders of magnitude in *δα*_1_ and keep the dilution rate constant for panel A, while panel B depicts the case with constant *δα*_1_ and a range of dilution rates *d*. The actual value of *d* can be discerned from the intersection with the 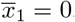 axis given by the single strain fixed point 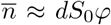. Only when the intersection of the isoclines occurs on the resource limited part of the population size isocline, we find a stable coexistence fixed point. As soon as this intersection moves (with changing parameters) through the transition to the antibiotic limited part of the isocline, this fixed point acquires complex eigenvalues, which become unstable and complex at the tip of the transition. Interestingly, at first this unstable complex fixed point becomes a stable limit cycle, with ever oscillating producer and non-producer populations. This is an example for a Neimark-Sacker bifurcation (Wiggins, 2003). However, moving the intersection further along the population size isocline, this limit cycle becomes unstable as well, and later the complex eigenvalues disappear, further down the antibiotic limited part, see Fig. 6.

**Figure 6.**
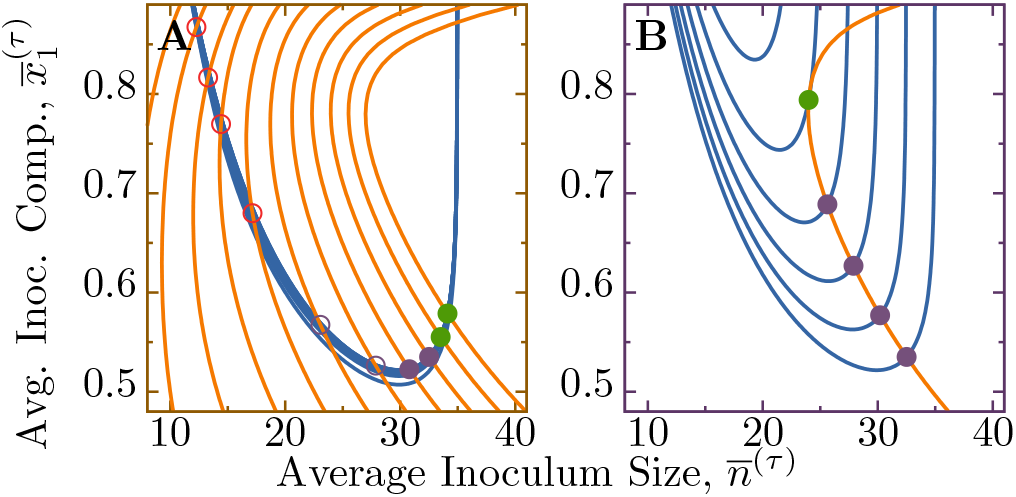
Zoom onto relevant parts of isoclines for antibiotic dynamics when changing the two main parameters. (A) Variation of growth rate difference, *δα*_1_ = 10^−1^ … 10^−3^, changes the composition isocline (orange lines). In contrast, this change in *δα*_1_ leaves the population size isocline (blue lines) almost unaffected: Isoclines for all choices of growth rate differences drawn on top of each other just appear with a larger stroke width, leaving individual curves indiscernible. As the composition isocline moves to larger inoculum sizes for decreasing *δα*_1_, it intersects the population size isocline at different positions. Intersections close to the transition between the two descriptions of the population size isocline (at minimal *x*_1_, see text) have complex eigenvalues, where an unstable, complex fixed point close to the minimum could actually hide a stable limit cycle. For panel (A), dilution rate is constant at *dS*_0_*φ* = 35. (B) Increasing the dilution rate shifts the resource limited part of the population size isocline. Populations start collapsing, when the inoculum size of the producing strain drops below the threshold value. For panel (B), growth rate difference is constant at *δα* = 6.31 · 10^−3^. Other parameters in both panels are *B*_0_/*μ* = 1.25 and *ρ*_1_/*α* = 5 · 10^−3^. Fig. 11 shows the stability of coexistence fixed points for a larger range of parameters.

In addition to these semi-analytical results, we argue that the stability of fixed point can be qualitatively checked by considering how the isoclines cut the phase plane, and which implications this has for the direction of a trajectory. In Fig. 5, shaded regions indicate inoculum sizes, where either the total inoculum size 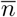, or the fraction 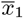 increases over one cycle. At a fixed point in the region of the resource limited part of the population size isocline, trajectories point towards the fixed point, which is thus stable. In contrast, the flow of trajectories indicates that a fixed point in the antibiotics limited part is unstable, as inoculum sizes will increase over cycles and move away from it.

### B. Iron extraction via siderophores

Another example of a public good is pyoverdine, which is an iron-chelating siderophore produced by several *Pseudomonas* species. Pyoverdine is a fluorescent extracellular molecule, which strongly binds otherwise (almost) unavailable iron and allows microbes to uptake the iron-pyoverdine complex via special transport proteins. Siderophores have often been considered to be a public good (Cordero *et al.*, 2012; Kümmerli and Brown, 2010; Lee *et al.*, 2016), but more recent experiments showed that this classification is highly dependent on details of environmental conditions (Julou *et al.*, 2013; Zhang and Rainey, 2013). Physiologically, enhanced iron-availability seems to increase the yield of cells (Clegg and Garland, 1971; Neilands, 1974), as populations grow to larger size with the same nutrients.

While the exact relation between yield and siderophore concentration is hard to specify, a few principles can guide our modeling. First, since cells require only minuscule quantities of iron and almost all experimental system will likely contain small traces of it, we assume that cells can maintain a minimal level of growth even without pyoverdine. Second, the effect of pyoverdine on yield saturates, with a maximum increase by a factor *σ*. In addition, pyoverdine is considered to be shared among all cells within a single deme, which corresponds to all strains expressing the appropriate transport proteins (Zhang and Rainey, 2013). Then, yield of all strains is affected simultaneously. Taking these considerations together, we propose

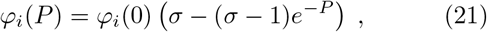
which is a simple exponential convergence towards a maximal value for increasing pyoverdine concentrations *P*. With this relation, yield is bounded by *φ_i_*(0) ≤ *φ_i_*(*P*) ≤ *σφ_i_*(0). Differences between strains are assumed as *φ_i_*(0) = (1 + *δφ_i_*)*φ*_0_, with *φ*_0_ a strain-independent constant. For the dynamics of pyoverdine, *P* (*t*), we assume,

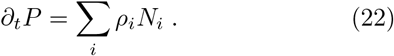

The rates *ρ_i_* are assumed to have all of the expression rate, excretion rate and the magnitude of their effect scaled in, such that *P* itself is a dimensionless quantity, that we can use in the exponential function of Eq. (21).

With these two additional relations, Eqs. (21) and (22), we can derive the expansion factor *ξ*. In these calculations we need to incorporate the effects of a time-dependent yield, as shown in the integrated resource use, Eq. (9). While details of the derivation are relegated to appendix C.1, our approximation is given by

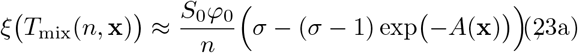

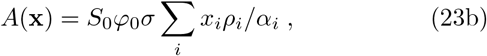
which is again ignoring the multiplicative corrections *G*(*n*, **x**) due to *δα_i_* and *δφ_i_* reported in Eq. (10). Corrections in *δα_i_* in addition to *G*(*n*, **x**) will likely occur from the pyoverdine dynamics itself, as Eq. (23) is essentially a zeroth order expansions in these differences. As in the case with antibiotics, the largest impact on final population sizes is given by the public good, as long as the increase in yield *σ* is large enough. Only when *σ* is small, *σ* − 1 ≪ 1, we need to consider these inherent differences between strains.

#### Analysis of isoclines

Having computed an approximation for the expansion factor *ξ* in Eq. (23), we can use this solution in the expressions derived for the isoclines. At first, we remark that saturation of the public goods dynamics occurs as long as *A*(**x**) ≫ 1. Then, the exponential term in Eq. (23) is negligible and we have *ξ* ≈ *S*_0_*φ*_0_*σ*/*n*. Accordingly, the single strain fixed point of a producer strain is 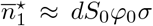. Since we assume that pyoverdine is not essential for growth, we also find a non-producer single strain fixed point at the already known position 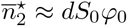. For two strains, the population size isocline then connects these two single strain fixed points: From the producer fixed point it extends down to lower producer fractions along a line given by Eq. (12b), with the replacement *φ* ↦ *φ*_0_*σ* to indicate saturated public good effects. At a fraction given by 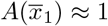, this saturation ends and final population sizes drop to their base level in absence of pyoverdine. Illustrations in Fig. 7 clearly show this behavior, where the (blue) population size isocline exhibits a sharp bend at small producer fractions. Similar to before, we can identify the dilution rate *d* as the main parameter influencing the position of this isocline.

**Figure 7.**
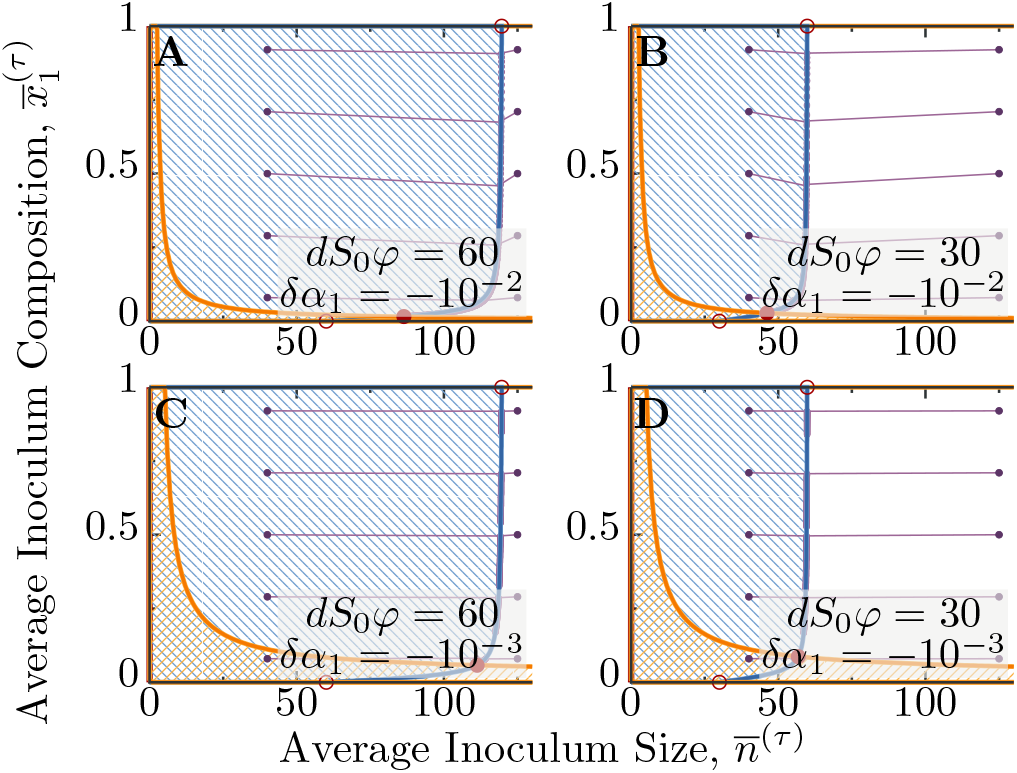
Dynamics over multiple cycles with production of pyoverdine that enhances iron-availability. Blue hatched areas indicate an increase of the average inoculum size 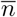 over one cycle, while orange hatched areas indicate an increase in the population composition 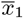. Strain 1 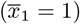 differs from Strain 2 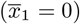 by a slower growth rate *δα*_1_ < 0 and a non-zero production of pyoverdine *ρ*_1_ > 0, *ρ*_2_ = 0. Purple dots connected by lines show exemplaric trajectories, that lead to coexistence fixed points for the chosen parameters. Only the first 50 cycles from each trajectory are shown. Adjustment of the average inoculum size 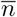 is fast, while the population composition 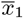 changes slow. Growth rate differences are chosen to be *δα*_1_ = −10^−2^ (A,B) and *δα*_1_ = −10^−3^ (C,D). Dilution rates (indicated by average inoculum sizes for growing populations) are *dS*_0_*φ* = 60 (A,C) and *dS*_0_*φ* = 30 (B,D). Other parameters are *σ* = 2 and *ρ*_1_/*α* = 10^−3^ for all panels.

For the composition isocline, we insert the expression for *ξ* into Eq. (13). Similar to the case with antibiotics, the main term contributing to a large positive covariance term is 〈*x*_1_(*ω* − 1)〉, which evaluates to

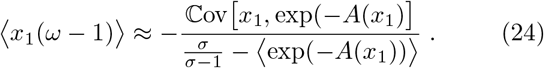

Interestingly, when numerically computing these averages over the Poisson distribution, we find that this expression decays exponentially close to 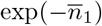, when *A*(1) ≫ 1. The factor *σ* only plays a role in the coefficient of this scaling. If *A*(1) ≪ 1, the dynamics effectively describes the original resource consumption, treated before, while for intermediate values, 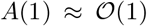, this expression starts to decay much slower. When focusing on the case with a significant effect of the public good, we can use the same reasoning that Eq. (24) needs to be balanced by terms of order 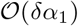: The shape of the isocline is approximately given by curves where expression (24) has a constant value, and thus follows along lines of 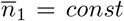. The position is again strongly determined by the growth rate difference *δα*_1_: For smaller *δα*_1_ the condition 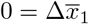 is met further out the tail of the exponential decay. Taking these considerations together, we find the scaling of the composition isocline as

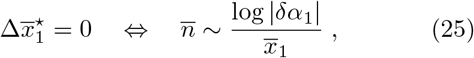
with a logarithmic dependence of *δα*_1_ on its position.

Coexistence can occur when isoclines intersect, which can again be achieved by changing the dilution rate. From checking the direction of trajectories around such fixed points, we find that coexistence is stable.

## V. Discussion

In this article, we investigated the dynamics of growing microbial populations that are repeatedly separated into compartmentalized demes on a long timescale *T*_mix_. Our main contribution is to provide a modelling framework that can encompass several different interaction types within microbial population and their environment. These interactions include growth on a shared resource, on top of which we treat explicit social traits like antibiotic degradation to allow all cells to grow, or pyoverdine expression, that enhances iron-availability for all of the population. We found that the spatio-temporal structure, together with the variance of initial conditions for the growth processes, allows costly traits that are helpful to the entire population to coexist stably over multiple cycles of seeding, growth and mixing.

The evolution of population composition is particularly insightful to derive the conditions of such coexistence. In repeated cycles of seeding, growth and mixing, we have shown that the change per cycle in the average inoculum composition is 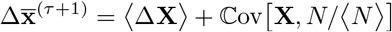 (see Eq. (4)). This describes two levels of selection whose origin lies in the ecological life cycle of populations: 〈Δ**X**〉 is the expected change of within-deme dynamics, usually dominated by fast growth.

The second term, 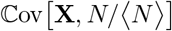, indicates correlations of the population composition with (relative) total size, and thus describes effects on a population level. If the population can generate a larger final size – due to reduction of antibiotics, enhanced iron-availability, or just more efficient resource conversion – this term can be large and positive, and thus can offset frequency losses from direct competition between strains. This frequency dynamics is also reminiscent of the Price equation (Okasha, 2006; Price, 1970). In our model, this connection to the Price equation arises naturally from the global mixing of the spatially distributed populations. Furthermore, Simpson-type effects are a special case for values of the terms in this frequency dynamics: The first term describing the expected within-deme change is negative and the average fraction inside demes decreases; but the covariance term is large enough to increase the average fraction in the entire system.

Similar models with spatial segregation have been considered, which also build on simple growth processes including public goods or other population level benefits. The work in (Cremer *et al.*, 2011, 2012; Melbinger *et al.*, 2010, 2015) is very related, where the authors deal with similar phenomena to those we considered: These include finite population sizes in segregated demes, fluctuations during the initial time of growth, and the second, long timescale on which all populations are mixed repeatedly. However, implementation details differ. Finiteness of populations is ensured by a (size-dependent) death rate, instead of explicit resource depletion. They identify fluctuations in cell numbers during initial stages of growth that amplify benefits to the whole population to be important for coexistence. Such fluctuations can leave traces in the population composition long after this initial time (Eggenberger and Pólya, 1923; Elhanati and Brenner, 2012). Our approach differs here in that such variability arises only from seeding; their implementation includes both stochastic seeding and stochastic growth dynamics. In another similar model (Ernebjerg and Kishony, 2011), the effects of fluctuations in the dilution rate *d* are examined: This can stabilize coexistence of similar strains over multiple cycles. Adding a lag-time before cells start to grow as third cellular parameter, next to growth rate and yield, also allows coexisting populations (Manhart *et al.*, 2018; Manhart and Shakhnovich, 2018), although restrictions on valid mixing times for coexistence of multiple strains exist.

One of the underlying assumptions going throughout most of our work, is that differences between different strains are small. This weak selection limit allowed to employ series expansions in both, *δα_i_* and *δφ_i_*, simplifying the dynamics. Such approximations are rather suited for organisms that are inherently similar, like microbes, while for communities of higher (more complex) organisms this might break down. For the resource consumption dynamics, however, the coexistence region in the differences *δα_i_* and *δφ_i_* extends to larger values compared to the two more elaborate models. In Fig. 4, these values even approach 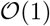. However, the required smallness of these parameters for antibiotics and pyoverdine dynamics can be used to our advantage: We reported that the population composition isocline shifts to larger inoculum sizes for decreasing *δα* (see Figs. 5 and 7, and Eqs. (20) and (25)). In turn, this shifts the (potentially stable) coexistence fixed point of the cycle dynamics to large inocula. We could also use this to our advantage: In experiments one could potentially deduce the growth differences between strains from this position of the fixed point – possibly with a resolution that is beyond contemporary fitness measurements. However, such measurements rely on being able to qualitatively predict the dynamical behavior within demes.

If growth rate differences are not too large, the population composition isocline exists off the boundaries, and thus the system is amenable to have a coexistence fixed points. Even with all other parameters constant, the dilution rate *d* can then provide an experimental knob to adjust the position of the population size isocline. Thus, diluting populations with different rates before they are seeded into demes is the easiest to generate an intersection of isoclines. For the simple resource consumption dynamics, we only find a restricted range of dilution rate that admit a coexistence fixed point, but for the two public good dynamics the shape of the composition isocline allows a much larger range of *d* that leads to coexistence. We argue, that also in natural populations, dilution rate will not be constant, and these fluctuations could at least temporarily allow coexistence of different populations.

Another relevant aspect to discuss is the importance of bottleneck sizes at the time of seeding. The magnitude of 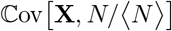 depends on how much both the final population sizes and population compositions vary, due to the inequality 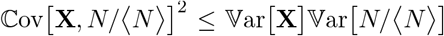. The averaging implicit in these expressions is computed over seeded inocula, whose spread scales as 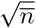. Consequently, a small inoculum size will have a twofold effect on the the overall covariance term, by affecting both factors in this bound: First, small 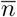 implies a larger spread over possible values, and thus can lead to a larger 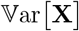. In addition, the enhanced granularity can generate more diverse initial conditions from which the growth process starts. Depending on the actual dynamics, this might lead to large differences in the final sizes, which are then related to the second factor in the bound, 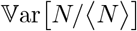. If inoculum sizes are large, the initial variability will play a smaller role in the deterministic within-deme dynamics: Final sizes do not have a large variation, and the covariance term plays only a minor role in the long term dynamics. These considerations can also be used for reasoning on ‘natural’ systems: In many contemporary experiments with microbes bottleneck sizes are large, and thus fast growth is often selected. In contrast, a majority of multicellular organisms go through a single cell bottleneck in their reproduction process (Grosberg and Strathmann, 1998). Limiting this bottleneck to only single cells clearly has disadvantages, but these could be offset by the gains due to the ‘population level’ effects (here essentially effects on the organism level) that can be encoded in large covariance due to cooperative processes during growth.

In addition, we may ask how strict the conditions need to be, which are required for our results to hold? We model a *perfect separation* of demes during growth, which are then instantly and globally mixed after a *constant* mixing time *T*_mix_. What would happen, if any of our modelling assumptions is subject to uncertainties? One way to weaken the assumption of separated demes is to investigate populations in continuous space with limited dispersal. In such a setting, it has been found that coexistence emerges on intermediary diffusion rates (Behar *et al.*, 2014; Oliveira *et al.*, 2014): Very fast diffusion makes the spatial dependence disappear altogether, while too slow diffusion leads to extinction of non-producers. Other mixing schemes than just a global pooling might alter the averaging, and thus also the eco-evolutionary dynamics of different strains. Recently, multiple groups tried to model the effects of specific interactions as a (directed) graph (Broom and Rychtář, 2008; Lieberman *et al.*, 2005; Taylor *et al.*, 2007), where – depending on topology of interactions – fixation of alleles could either be enhanced or hindered. While most of these analyses treat each of the interacting nodes as individuals, we expect that if each of them encompasses its own population dynamics on a multilevel approach like ours, the dynamics could be again skewed in either direction. Finding answers to this issues would be one possible avenue for future work.

In summary, we analyzed models of social interactions of spatially distributed microbial populations. Our results showed that coexistence – also of costly social traits – can be traced back to the simple ecological mechanisms of a second timescale and spatially distributed populations. The dynamics of these traits can be described by an expression akin to the Price equation, which allowed reasoning within a framework that generalizes several previously published modeling approaches. Moreover, we also expect other collective dynamics to show similar behavior (Payne *et al.*, 2018), when they are subject to similar spatio-temporal structuring of the environment.

## Acknowledgments

The authors thank B. Altaner, D. Hofmann, F. Schüßler and L. Susman, in addition to other members of the NBR Labs for discussions and comments on the manuscript. This work was supported by HFSP Grant RGP0010/2015.

## Appendix A: Growth dynamics of resource consumption

In the following, we present more technical details for the resource consumption dynamics to justify some of the expressions stated in the main text. The extensions to the interactions including either antibiotic reduction or enhanced iron-availability from pyoverdine are treated in their own sections B and C below.

### 1. Growth and stability of a single strain

At first, we treat a simplification of the overall dynamics: A single strain, with constant growth rate *α* and constant yield *φ*, feeding on a finite resource, and without any other environmental interactions. This is most likely the most basic model that still fits our framework, which allows to gain insight via analytic solutions.

The within-deme dynamics in this case is

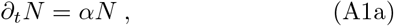

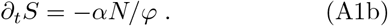

Clearly, we find *N*(*t*; *n*) = *n* exp(*αt*) as long as cells are growing. This expression can be inserted into the resource dynamics, which in turn can be integrated analytically from *S*(0) = *S*_0_ to *S*(*T*_depl_) = 0: we obtain 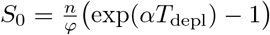, which can be inverted to find explicit depletion time *T*_depl_ as a function of *n* and the other parameters. Assuming that growth ceases upon resource depletion, *α*(*t* > *T*_depl_) = 0, we find the full solution for population size to be

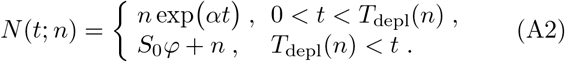

For the dynamics over multiple cycles, the time is constant at *T*_mix_, and Eq. (A3) is assumed to be an function only of inoculum size *n*. Then, depending on the number of cells present at the beginning, resources are either used up and the second expression is relevant, or the population is still in its growth phase and we need to use the first expression. These situations are depicted in Fig. 8 as dashed lines. Over multiple cycles the mapping takes the form

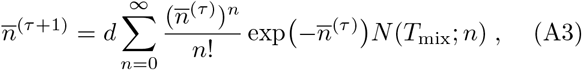
where the average number of grown cells is multiplied with the dilution rate *d* to account for the seeding into new demes. The Poisson distribution for seeding probabilities used here smoothly interpolates the two branches of the population size in Eq. (A2).

One of the features observable from the mapping of Eq. (A3) (and Fig. 8) is that resources need to be depleted for the mapping to have a stable fixed point. A stable fixed point of this mapping can only exist in the upper, stationary branch of the solution. If cells would still be growing at the time of mixing *T*_mix_, the lower branch of the solution is used for the mapping, which does not support a fixed point. However, this is just a necessary condition, not yet sufficient. If the dilution rate *d* is too small we do not expect a fixed point either.

**Figure 8.**
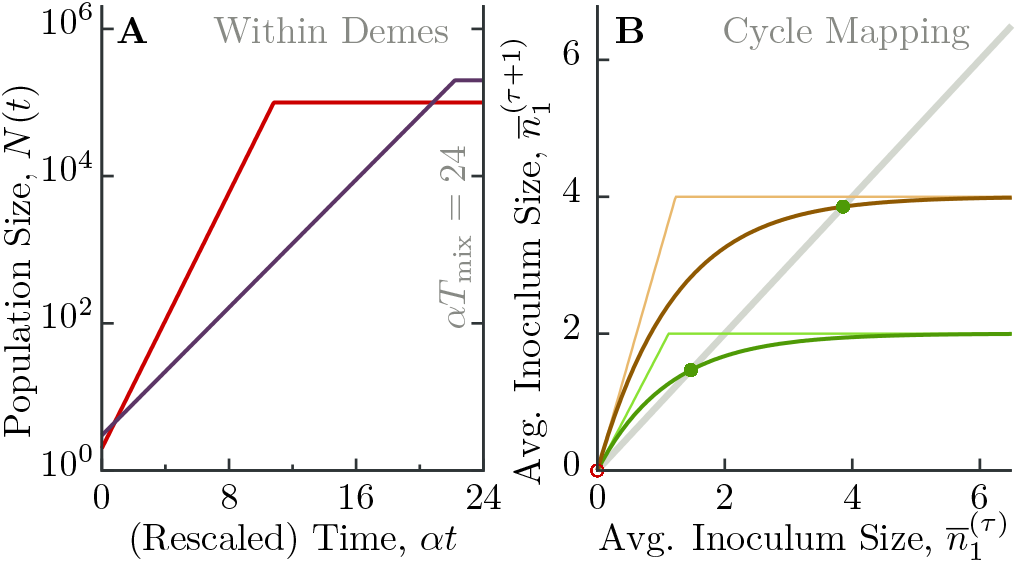
Dynamics of a single strain with only resource consumption. (A) Trajectories over time for two different inoculum sizes *n* and growth rates *α*_1_. The population grows exponentially and then stays at constant numbers. (B) The inoculum size for a single is averaged with a Poisson distribution over all possible inoculum sizes. This smears the sharp transition from starting with enough cells that deplete all nutrients to inoculum sizes that are still in the exponential growth phase at *T*_mix_ (dashed lines). As explained in the text, existence of a fixed point depends on *dN*(*T*_mix_;1) > 1, and thus how many cells in the inoculum If this condition is not met, the population will be washed out over multiple cycles.

In order to compute the stability of a fixed point, we introduce an auxiliary calculation that will help in this respect. Specifically, taking a derivative with respect to the parameter of a Poisson distribution, we have

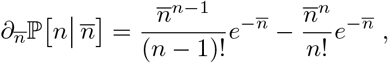
which leads to a shift in this summation index, 
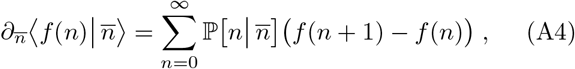
because we are only concerned with functions that exhibit the property *f*(0) = 0. This property indicates the obvious fact that if the inoculum is empty, nothing will grow. This expression generalizes straightforward to multidimensional variables, 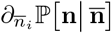, where the shift happens only for strain *i*.

With this calculation in place, we can compute the slope of the cycle mapping for a single strain at the origin. This allows to check if extinction is a stable or unstable fixed point, where the latter implies that another stable non-zero fixed point exists. Formally, we obtain

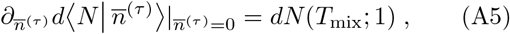
as for 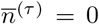 the Poisson distribution collapses and only the first term in the auxiliary derivation above survives with an inoculum of 1. The extinction fixed point is unstable if it is larger than 1, and stable otherwise. When nutrients are not yet depleted at this time, the condition (A5) evaluates to *d* exp(*αT*_mix_), which needs to be increased above 1 to find a fixed point. Assuming that the growth rate *α* is not changed as easily, increasing the mixing time *T*_mix_ has the larger impact, which clearly makes sense: if the population has more time it can deplete nutrients. If nutrients are already depleted, but the slope is still below 1, we find that 1 > *dS*_0_*φ*. Thus, an average inoculum size of less than a single cell is also not stable. In these cases, the population will be washed out over consecutive cycles.

To proceed, we assume that mixing time is large enough that even a single cell uses up all nutrients. In this case, we can write the final number of cells as *N*(*T*_depl_; *n*) = (1 − *δ_n_*_0_)(*S*_0_*φ* + *n*), where *δ_n_*_0_ is a Kronecker-delta to indicate the fact that nothing can grow from an inoculum size *n* = 0. Then, the sum in Eq. (A3) can be evaluated, and we find the dynamics 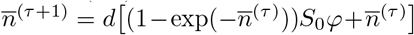. From this expression, we can compute the fixed point 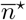,

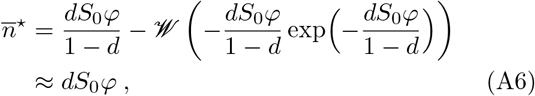
where 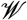 was the Lambert-W function. The correction indicated by 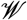 becomes only large when *dS*_0_*φ* is close to 1, at which it is approximated by 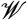 (.) ≈ exp(−*dS*_0_*φ* +1). As we argued above, this full expression for fixed point also demonstrates that an average inoculum size below a single cell will not be stable, as the expression evaluates to zero for *dS*_0_*φ* < 1. For a large parameter combination of *dS*_0_*φ*, the Lambert-W function is almost linear in its argument, such that 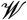 (−*y* exp(−*y*)) ≈ *y* exp(−*y*), and we obtain an exponential correction for the single strain fixed point, 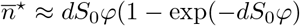. This correction can explain for instance, why the population size isocline in Fig. 3 is shifted slightly off the value of the linear approximation in Eq. (A6).

Stability of this non-zero fixed point can be checked by

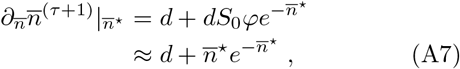
which is smaller than 1 for any realistic value of *d*, since the term involving the exponential is bounded, ≤ 1/*e*. Thus, the fixed point given by Eq. (A6) will be stable, if it exists.

### 2. Coexistence of multiple strains

After establishing the value of the single strain fixed point in Eq. (A6), we turn to the case with multiple strains. In the main text, existence and stability has been explained using isoclines in total population size and the population composition. Essentially, this allowed to find similarities and differences when comparing with the two examples for public good interactions. Here, however, we derive a few additional results on the resource consumption dynamics. These calculations benefit from a description in absolute numbers 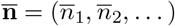 for each of the strains, and not use the non-linear transformation of computing fractions 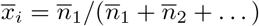.

One way to define coexistence in this system of multiple strains, is when *all* single strain fixed points are unstable with respect to invasion by other strains. Formally, this happens when the Jacobian evaluated at these single strain fixed points features eigenvalues larger than 1. We already derived that a strain that does not use up all resources will be washed out over cycles, which also holds for multiple strains: For the simple resource consumption for the within-deme dynamics, each strain needs also to be stable alone, if it should be part of a coexisting collective. Then, if *all* of these fixed points are unstable with respect to invasion by other strains, the system will have an additional fixed point that indicates coexistence, see Fig. 3. Entries of the Jacobian 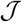 at these single strain fixed points 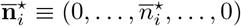 can be computed by

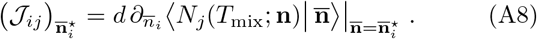

For the following, we assume that the resident strain has index 1. We already computed 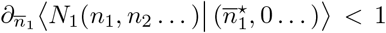 in Eq. (A7), which just indicates that the resident strain is stable on its own. In addition, other relevant computation include how average final sizes change when changing any of the inocula 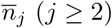. Clearly, as no strain other than 1 is present, final sizes of strain *i* stay at zero upon changing the inoculum of other strains 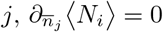, as long as both *i* and *j* are different, and neither is the resident strain. Thus, the Jacobian is in general a very sparse matrix, which only has entries on the diagonal and in the first line. This structure of the Jacobian implies that all entries on the diagonal are already the eigenvalues, as it is a upper triangular matrix. For the entries on the diagonal itself, we compute

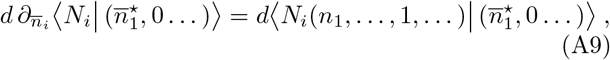
for *i* ≥ 2, where we used the auxiliary calculation in Eq. (A4) from above. The average is taken at the single strain fixed point of strain 1, and the inoculum includes *exactly* a single cell of strain *i*. Thus, if this single cell of strain *i* grows to more than one cell (on average) in the inoculum of the next cycle (eigenvalue larger than 1), the single strain fixed point of strain 1 is unstable with respect to invasion of strain *i*. As all single strain fixed points need to be unstable with respect to each other strain – this leads to *k*(*k* − 1) pairwise conditions for *k* strains. However, any subset of strains that fulfills these pairwise conditions can exhibit coexistence.

The coexistence regions depicted in Fig. 4 are obtained by evaluating these conditions for two strains numerically. Then, only the two conditions 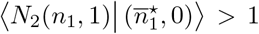 and 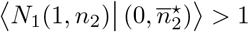 need to be fulfilled.

Note, however, that these set of conditions (A9) is not yet constructive: when one set of strains with their growth parameters *α_i_* and *φ_i_* is given, one can check if they would be able to coexist. However, it does not show how to choose these parameters to get coexistence between an arbitrary number of strains. Numerical estimates indicate the the possible range of values for these two parameters decreases more and more with each additional strain. In particular, in (Ernebjerg and Kishony, 2011) the authors chose values on a one-dimensional curve in *δα* and *δφ* to find coexistence between multiple strains.

### 3. Evaluating relations for the composition isocline

In the main text, we reported an expansion of 〈Δ*X_i_*〉 and 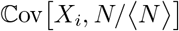 in small growth rate differences and yield differences, see Eq. (14). Here, we present the steps to obtain the scaling announced in Eq. (15) via evaluating these expressions. For two strains, we have

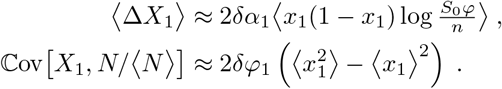

The two independent Poisson distributions for the inoculum sizes *n*_1_ and *n*_2_ transform into a Poisson distribution for the total size *n* = *n*_1_ + *n*_2_ and a binomial distribution for *n*_1_. This transformation has the effect, that the sum over *n* is started at 1, as without any inoculum the population will not grow. As a next step, we need to evaluate the following terms occurring in the expressions above,

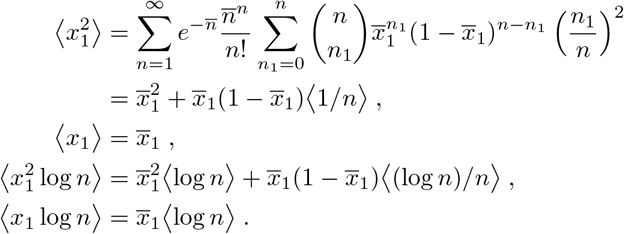

Using these, the balance for all terms in the expansion of the composition isocline is given by

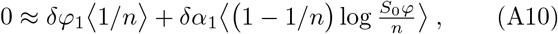
from which we extract the approximation reported in the main text,

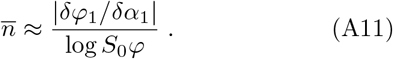

Numerical observations suggest that this scaling also contains a coefficient that is slightly smaller than 2. However, we are mostly interested in how this position of the composition isocline scales with different parameters in the model.

## Appendix B: Growth dynamics with antibiotics

Originally, we stated the coupling to environmental processes as an ODE in addition to Eqs. (5). However, our formalism uses expansion factors *ξ*(*T*_mix_(*n*, **x**)) to represent this interaction of strains with their environment. This section presents the derivation of how to obtain this expansion factor *ξ* and a few additional results on the dynamics of antibiotic reduction.

### 1. From dynamical effects of interactions within demes to expansion factors

The effect of an antibiotic concentration *B* on growth rate is defined via Eq. (16), which describes a sigmoidal, decreasing curve for increasing concentrations *B*. In this expression, the antibiotic concentration is measured in multiples of the ‘MIC’ *μ*, which is the concentration where the population switches from overall death to overall growth. The set of relevant coupled differential equations to analyze is given by

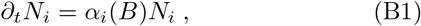

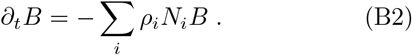

In the following, we present an approximation scheme that captures the essence of the dynamics. Somewhere else we show that our approximation is actually very robust against changes of molecular details, and can describe several classes of antibiotic resistance (manuscript in preparation).

The first step in solving the dynamics is defining a logarithmic antibiotic concentration *K* = log(*B/μ*). Often, only the magnitude of this concentration is important, and not its exact value. Using this logarithmic concentration, we can approximate the effect of antibiotic concentration on the growth-/death-rate of populations as 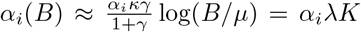. This allows to write the second equation for the reduction of antibiotics as *∂_t_K* = − *i ρ_i_N_i_*. Furthermore, we assume that cells die with a rate given by the initial antibiotic concentration, *N_i_*(*t*) = *n_i_* exp(−*α_i_λK*_0_*t*. Then, we can integrate the dynamics for antibiotic decay, which gives

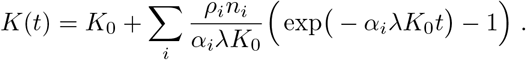

We define a time *T*_1_ implicitly via *N*(*T*_1_) = 1, ie. when the exponential decay of the population reaches a single cell, 1 = *n_i_* exp(−*α_i_λK*_0_*T*_1_). This time *T*_1_ can be inserted into the dynamics of the logarithmic antibiotic concentration *K*(*T*_1_), to check if this concentration decayed already below 0 (population survived), or still exceeds 0 (population will go extinct):

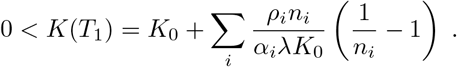

As long as the inoculum sizes are large enough, 1/*n_i_* 1, we find the inequality

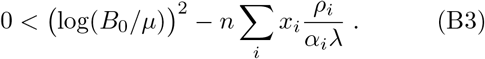

This condition, Eq. (B3), indicates that the microbial population reduces the antibiotic concentration fast enough that the overall population again exhibits a positive growth rate before going extinct. From there on, the surviving – and growing – population will degrade the remaining antibiotic fast. We assume that the mixing time *T*_mix_ is long enough, that the population will use up all remaining nutrients. As the dying cells during this initial phase are likely few compared to the final size, these final sizes are close to the population sizes obtained from the simple resource consumption model. Consequently, the expansion factor can be written as

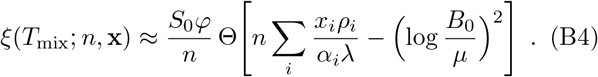

The Heaviside-Theta function Θ indicates exactly the condition (B3) above, where the expansion factor evaluates to zero, if all cells die, and is approximated by the first order term *S*_0_*φ/n* of Eq. (10) when they survive.

For a single strain, we illustrate how this resulting expansion factor generates the mapping of inoculum sizes over cycles in Fig. 9. The sigmoidal curve emerges from smearing out the sharp cutoff between surviving and extinct populations. Exactly this sigmoidal shape implies that either two or no (coexistence) fixed point occurs in the mapping.

**Figure 9.**
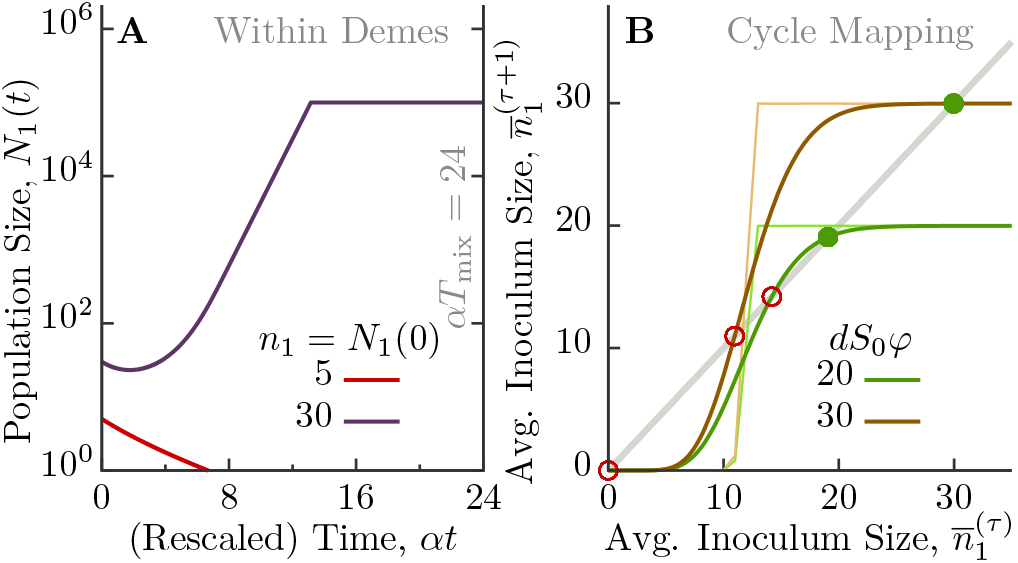
Dynamics of antibiotic reduction for only a single producer strain. (A) Within demes, the initial population *n*_1_ either manages to reduce antibiotics in time (purple trajectory), or fails and goes extinct (red trajectory). (B) In the mapping of average inoculum sizes over cycles, this threshold in inoculum size between survival and extinction generates a step-like increase of the cycle map. Dashed lines indicate the expected inoculum size if the dynamics within demes would start with an inoculum of *n*_1_, while solid lines indicate the mapping for the average population size *n*_1_. The latter includes a range of possible inoculum sizes sampled from a Poisson distribution, which smears out the hard cutoff between survival and extinction. Depending on dilution rate *d*, either two or no (non-zero) fixed points exist. Parameters are *S*_0_*φ* = 10^5^, *ρ*_1_/*α* = 5 · 10^−3^, *B*_0_/*μ* = 1.25.

### 2. Additional parameter dependence of the cycle mapping

In the main text, we mostly focused on how isoclines change upon varying the two main parameters *δα*_1_ and *d*. As we explained, each of them changes one of the two isoclines (population size or population composition, respectively), while leaving the other isocline almost unaffected. However, the models for antibiotic reduction also features additional parameters, whose effects we only report here.

The dynamics of antibiotic reduction clearly depends on how much antibiotic *B*_0_ is present at the beginning of the growth phase and how fast antibiotics is reduced. From the threshold for growth/no-growth, as stated in Eq. (B3),

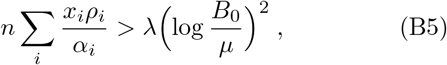
we see that both *ρ_i_*/*α_i_λ* and *B*_0_/*μ* give additional important parameter combinations. Fig. 10 explores the effects of changing either of those. In general, as long as *B*_0_/*μ >* 1, the effect of changing either production rate or initial antibiotic concentration only changes the threshold inoculum size for producing strains that would survive. However, when *B*_0_/*μ* < 1 the non-producing strain will be able to survive on its own, although it might grow significantly slower in demes where antibiotic concentration is not reduced. Due to this survival, the part of the composition isocline that indicated the extinction of the non-producing strain (asymptotic approach towards the 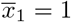 axis) disappears, and the composition isocline directly intersects with the producer axis of the phase plane, see Fig. 10AB.

**Figure 10.**
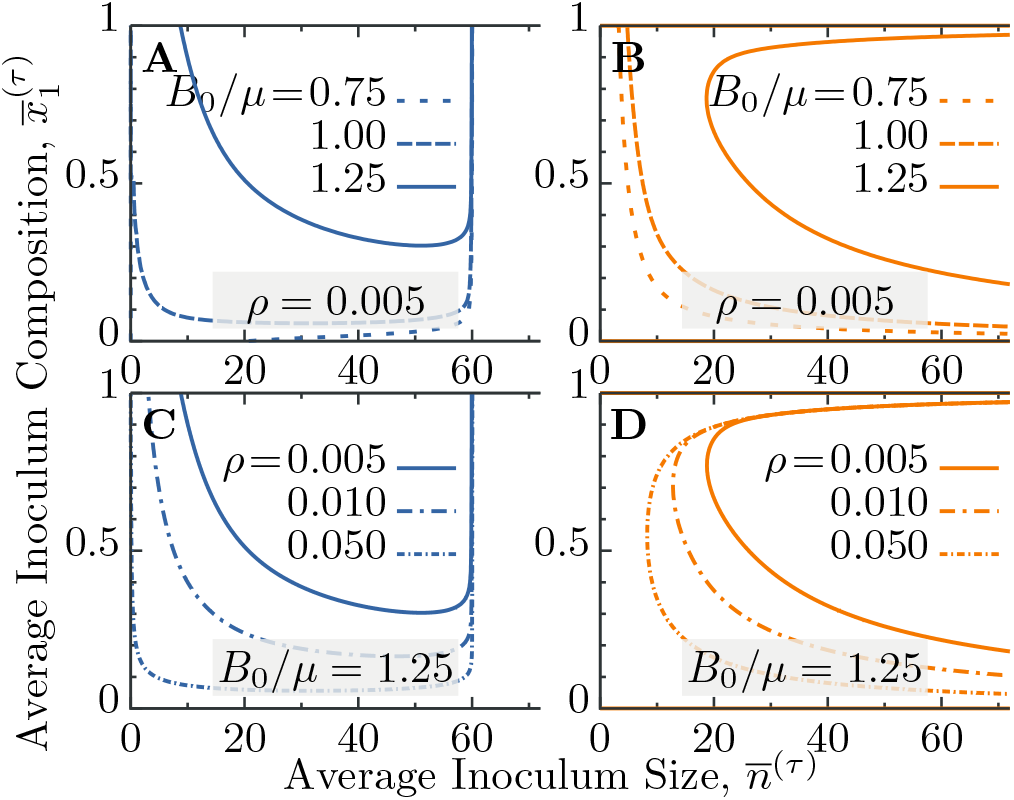
Influence of other parameters on the shape of isoclines in the dynamics with antibiotics. Panels show the isoclines for (A) average inoculum size *n* and (B) average inoculum composition 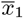, when varying the initial antibiotic concentration *B*_0_/*μ*. In (A), from top to bottom these initial values are *B*_0_/*μ* = 1.25, 1.00, 0.75. In (B), the innermost curve indicates the highest antibiotic concentration, while the outermost curve the lowest concentration. Panels (C) and (D) show the effect of changing the production rate *ρ*_1_ of the enzyme. Parameters *dS*_0_*φ* = 60 and *δα*_1_ = −10^−2^ are constant in all panels.

### 3. Stability of the coexistence fixed point

We can check existence and stability of the coexistence fixed point numerically. In the main text, we reported that this coexistence fixed point transitions from stable to complex stable to complex unstable (briefly with a stable limit cycle) and finally to unstable, when shifting the intersection of the isoclines along the population size isocline by altering the growth rate difference. Fig. 11 shows these results for a range of the two main parameters dilution rate *d* and growth rate difference *δα*_1_ of two strains. There, we specifically focus only on the (potentially stable) coexistence fixed point, and not on any other of the fixed points of the cycle mapping. Thus, this stability analysis does not specify the exact position of this fixed point, but its stability can give already an qualitative indication on which branch of the population size isocline it occurs.

**Figure 11.**
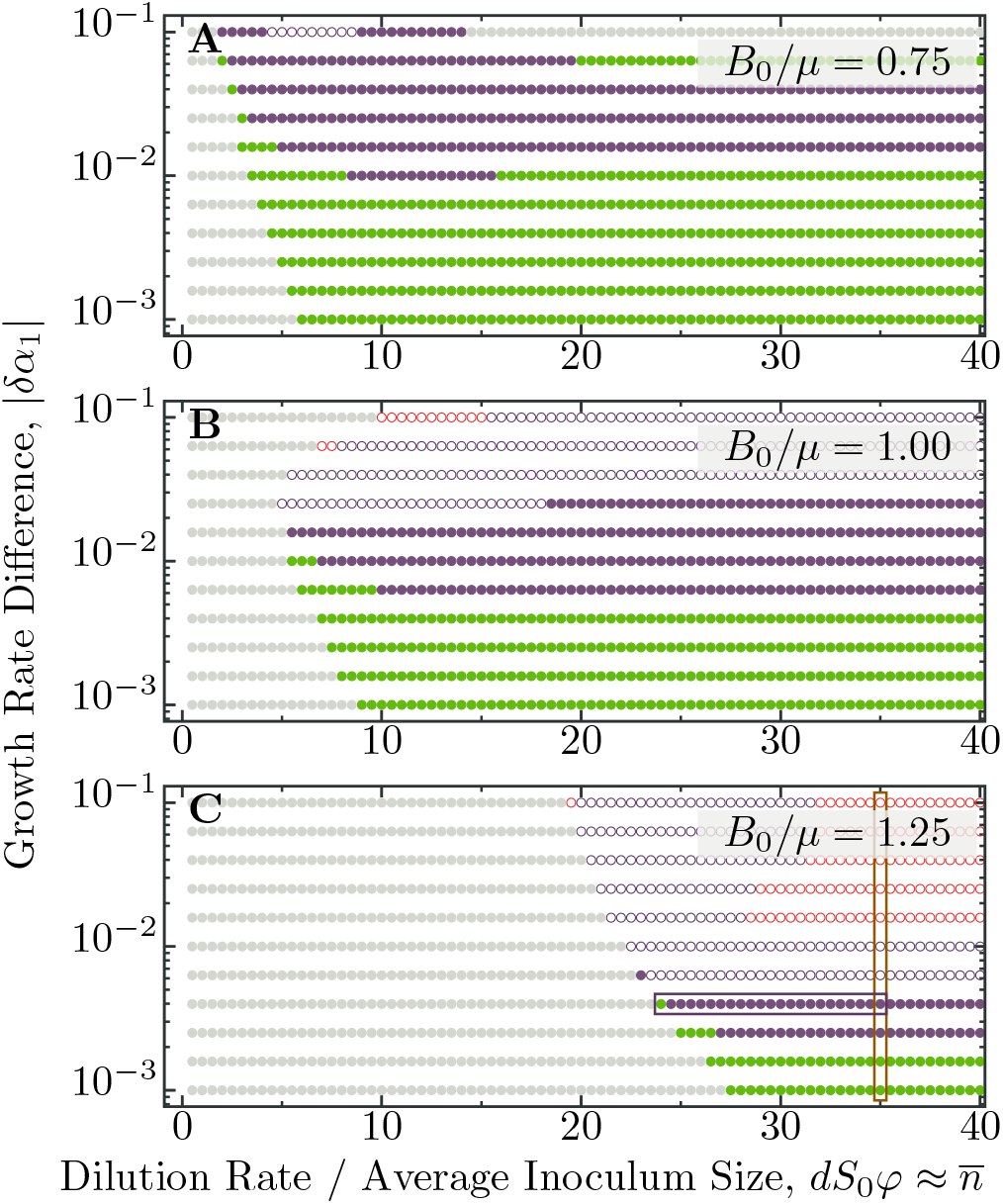
Stability of fixed points within-deme dynamics involving antibiotic hazards. Displayed are color-coded stability properties of the coexistence fixed point, as long as it exists (gray symbols indicate it does not). Specifically, the colors indicate stable (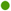), stable with complex eigenvalues (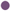), unstable with complex eigenvalues (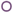) and unstable (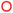) fixed points. In the complex unstable regime, close to the boundary to stability, we also find stable limit cycles, which are not captured in the linear stability analysis of computing eigenvalues of the Jacobian. These limit cycles appear through a Neimark-Sacker bifurcation (Wiggins, 2003). The change in shape and position for the two isoclines of inoculum size and inoculum composition are shown in more detail in Fig. 6, where purple and brown boxes in panel (C) correspond to the similar colored borders.

## Appendix C: Growth dynamics with pyoverdine

In the main section, we only reported the result of our approximation for the expansion factor *ξ*. Here, we provide the steps in the derivation of this approximation, which is effectively zeroth order in the growth rate differences *δα_i_*.

### 1. From dynamical effects of interactions within demes to expansion factors

Pyoverdine changes yield *φ* on a population level due to enhanced availability of iron (see Eq. (21)),

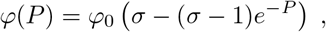
where the production of pyoverdine *P* is given by *∂_t_P* = Σ*_i_ ρ_i_N_i_*. In order to compute its effects on the expansion factor *ξ*, we need to evaluate the full, time-dependent resource-use equation (9). This equation requires to evaluate an integral that contains a term how yield changes over time, *∂_t_φ*. Consequently, the dynamics has to be combined with the production dynamics of pyoverdine, Eq. (22).

To this end, note that both time-evolutions of *S*(*t*) and *P* (*t*) are linear in *N*(*t*), which allows to combine their two dynamics as

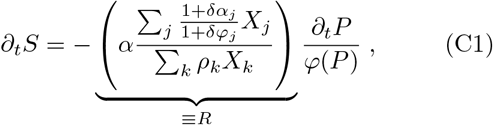
where in anticipation of the results we wrote the dynamics of the pyoverdine concentration *P* together with yield *φ*(*P*), which is influenced by this concentration. The population composition *X_i_*(*t*) changes only slowly in time, thus we assume that *X_i_*(*t*) ≈ *x_i_* for the purpose of the ensuing approximations, and collect all these dependencies in a constant *R*. Then, we can integrate this equation from initial conditions *S*(0) = *S*_0_ and *P* (0) = 0 up to a time *t* to get the relation between *S*(*t*) and *P* (*t*),

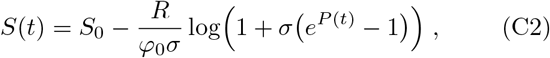
which is an almost linear expression of the form *S*(*P*) ≈ 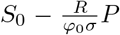. As a next step, we take a derivative with respect to time on *φ*(*P*), Eq. (21), to obtain

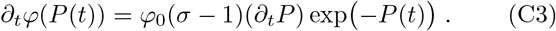

With these two expressions, we can evaluate left hand side in Eq. (9),

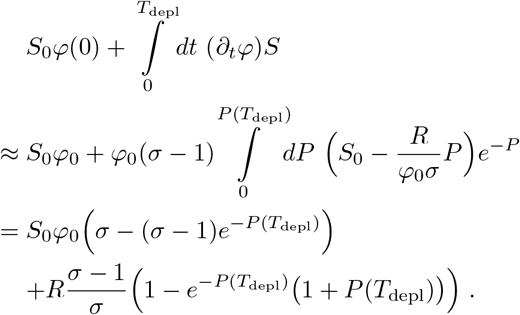

The first line in this solution indicates the expected yield at the end of the growth phase. The expression in the second line denotes the correction due to the fact that the concentration of *P* is increasing during this growth phase, and did not yet start at the final value.

Having this expression, we can simplify further, and neglect the second line to obtain the main scaling of the expansion factor. To this end, we define the abbreviations *ξ*_0_ = *S*_0_*φ*_0_*σ*/*n*, and furthermore 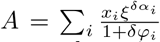. With these, the full resource use equation can be written as

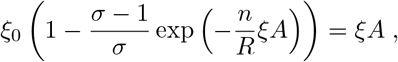
which we need to solve for *ξ*. Note that *R* also contains a logarithmic dependency on *ξ*, which does not influence the overall scaling. The term *A* generates the corrections in *δα_i_* and *δφ_i_* reported in Eq. (10): Here we see that they are multiplicative factors, which is the reason that we can solve for *ξA* first. We find that

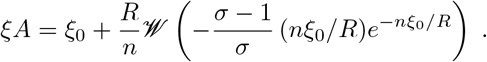

For small enough arguments the negative branch of the Lambert-W function is linear in this argument, which in this expression is usually sufficiently suppressed by the exponential term. Thus, we can simply leave out 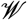 of this expression to obtain the final scaling of the expansion factor for pyoverdine production,

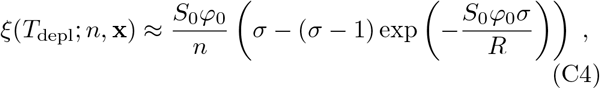
which assumes *A* ≈ 1. Deviations from unity generate the corrections *G*, as shown in Eq. (10).

### 2. Other parameter dependence

The main text only explored the effects of changing growth rate differences. Similarly, the dynamics of pyoverdine expression to increase iron-availability exhibits also a production rate dependence, *ρ*. Moreover, here the factor *σ* by which yield is increased plays another important role. Fig. 12 explores the effects on isoclines if either of these are changed.

**Figure 12.**
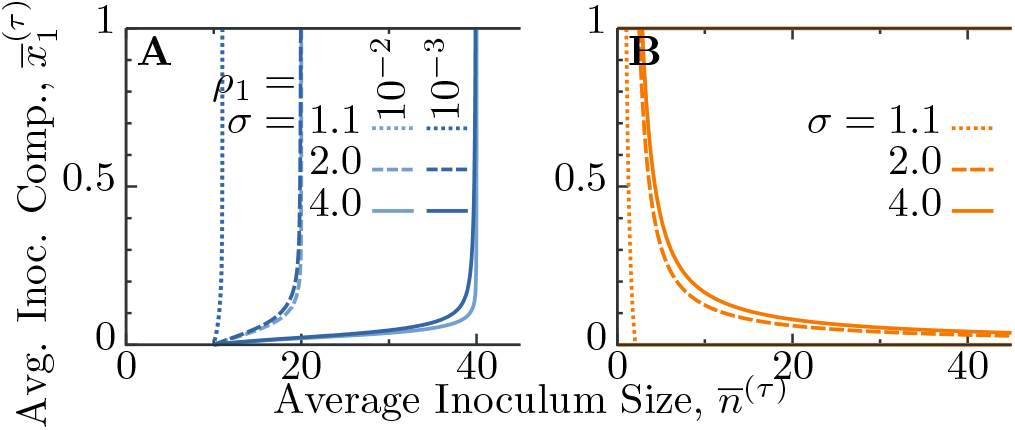
Influence of other parameters on the isoclines in the dynamics with pyoverdine. Panels show the isoclines for (A) average inoculum size *n* and (B) average inoculum composition *x*_1_, upon variation of the factor *σ* that increases yield due to presence of pyoverdine. In panel (A) we additionally illustrate the influence of the production rate *ρ*_1_ as different shades of blue. This production rate does not significantly change the isoclines in (B). Parameters *dS*_0_*φ* = 10, *δα*_1_ = −10^−2^ are constant for all plots.

## Appendix D: Numerical methods

Several results in our manuscript have only been obtained numerically, although we tried to explain these results with scaling arguments. Here, we briefly present numerical methods used throughout our work.

In general, the two levels of selection of growth within demes and the mixing cycles are also reflected in our code. We first solve the within-deme dynamics, on a grid of all possible initial conditions (*n*_1_*,n*_2_). These initial conditions are chosen up to values *n_i,_*_max_, such that the probability of finding this combination in the Poisson distribution are tiny and negligible. Often, we use the condition 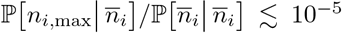 to determine these maximal inoculum sizes. For most of the results shown in Figs. 5 and 7 this maximal inoculum size is roughly *n_i,_*_max_ = 200, which is the value we use in our figures. Solutions to within-deme dynamics are obtained by a standard Runge-Kutta 4th order integration scheme up to the time *T*_mix_.

After final population sizes *N_i_*(*T*_mix_; *n*_1_*,n*_2_) are computed, they are stored as lookup-table. Cycles of growth, mixing and reseeding are realized by computing the sums

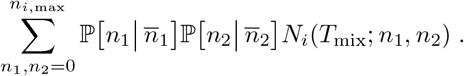

Fixed points of the mapping (e.g. Fig. 11) are obtained via iteration of a multi-dimensional Newton-Raphson scheme, which requires usually only a few steps.

Code is available at https://github.com/lukasgeyrhofer/mixingcycles under the CC0 license.

